# Unveiling *Crocosphaera* responses to phosphorus depletion: insights from genome analysis and functional characterization

**DOI:** 10.1101/2024.10.09.617355

**Authors:** Chloé Caille, Sophie Rabouille, Eva Ortega-Retuerta, Yann Denis, Olivier Crispi, Barbara Marie, Mireille Pujo-Pay, Vladimir Daric, Emmanuel Talla, Amel Latifi

## Abstract

Unicellular, nitrogen-fixing cyanobacteria (UCYN) thrive and support primary production in oligotrophic oceans, playing a significant role in the marine nitrogen cycle. *Crocosphaera* sp, a model for studying marine nitrogen fixation, is adapted to low phosphate (P_i_) conditions. Yet, how *Crocosphaera* copes with P_i_ depletion is rather poorly understood. We present genomics analysis of P_i_ stress-responsive genes in this genus, encompassing six *C. watsonii* genomes and two strains isolated in coastal environments, *C. subtropica* and *C. chwakensis*. We identified genes involved in P_i_ signaling, uptake, and dissolved organic phosphorus (DOP) hydrolysis. Results showed different genetic potentials to cope with P_i_ scarcity between the *Crocosphaera* strains. Physiological monitoring of cultures of *C. watsonii* WH8501 exposed to P_i_ depletion highlighted a capacity to divide several times and survive for a few more days, albeit with a skewed C:N:P stoichiometry. Upon addition of DOP, cultures efficiently recovered to a growth rate and cell composition equivalent to those observed under favorable conditions. The concomitant transcription analysis revealed diel expression patterns of P_i_-related genes and endogenous clock genes, suggesting a possible circadian regulation. Our data deepen our understanding of the growth strategies *Crocosphaera* employs in P_i_-limited environments, offering broader insights into microbial resilience in marine ecosystems.

## Introduction

Unicellular, nitrogen-fixing cyanobacteria (UCYN) are prevalent in the tropics and subtropics areas of the open ocean, playing a significant role in the marine nitrogen cycle (1). Nitrogen (N) fixation is the primary mechanism introducing new N into the open ocean, supporting up to half of the new primary production (2) (3). This underscores the critical importance of diazotrophic cyanobacteria in these oligotrophic regions. The genus *Crocosphaera* is a cultivated representative of open-ocean, photo-autotrophic UCYNs. It is a key model organism for studying N fixation and how this process varies with environmental factors. Based on the phylogenetic distribution of the *nifH* gene, the *Crocosphaera* genus has been divided into two clades: the UCYN-B clade, which includes the species *C. watsonii* (4) (5), further subdivided into small and large strains (6). The UCYN-C clade includes *C. subtropica* and *C. chwakensis*, two strains isolated in coastal environments and formerly classified under *Cyanothece* (7). The growth of diazotrophic cyanobacteria can be limited by the availability of nutrients, primarily iron and phosphorus. Nevertheless, *Crocosphaera* strains are widely distributed in oligotrophic regions with low phosphorus (P) levels (1). For instance, *in situ* observations indicated that diazotrophic cyanobacteria are present the Southwest Pacific (8) and the Northern tropical Atlantic Oceans (9). Their ecological distribution suggests that *Crocosphaera* strains have developed adaptive mechanisms to cope with P_i_ depletion.

Bacterial strategies to adapt to phosphate (P_i_) limitation are diverse, including the replacement of phospholipids with sulfolipids, the use of high-affinity P_i_ transporters (10), and the scavenging of dissolved organic phosphorus (DOP) sources through hydrolytic enzymes (4) (11) (12). Most of these mechanisms are induced in response to P_i_ depletion, as perceived by the two-component regulatory system of the PhoB-PhoR family (13). Current knowledge on how *Crocosphaera* copes with P_i_ depletion is primarily derived from studies on *C. watsonii*. For example, alkaline phosphatase activity was measured in *C. watsonii* P_i_-depleted cultures and this strain can use dissolved organic phosphorus (DOP) as its sole phosphorus source, including phosphomonoesters and phosphodiesters (14). However, the transcription of none of the three genes annotated as potentially encoding alkaline phosphatase enzyme was induced in response to P_i_ deficiency (15) (16). When extracellular P_i_ is abundant, *C. watsonii* employs the constitutively expressed phosphate inorganic transport (Pit) system to import phosphorus into the cell (15). Under P_i_ limiting conditions, a high-affinity phosphate-specific transport (Pst) system, which uses ATP-mediated transport, is activated, and the high-affinity phosphate-binding protein (PstS) is upregulated to enhance P_i_ uptake (16) (17). Comparative genome analysis of six unicellular *C. watsonii* strains belonging to the two morphological phenotypes (small and large cells) revealed a variation in the copy number of genes involved in the transport and use of P_i_ (6). Whole-genome transcription analysis of *C. watsonii* cultures showed that 47.4% of the genes exhibited a diel expression pattern (18). In P_i_-depleted cultures, the *pstS* gene also displayed a diel expression pattern (16), indicating that adaptive mechanisms to cope with P_i_ scarcity might follow diel regulation, likely by the circadian KaiABC clock (19).

The results summarized above point to results specific to *C. watsonii*, while a comprehensive comparative overview of the genetic potential of the genus *Crocosphaera* to cope with P_i_ limitation is lacking. Here, we present a dual approach to the genetic and physiological response to P_i_ limitation. First, to compare the genetic potential of the different *Crocosphaera* strains, we conducted a functional genomics analysis of P_i_ depletion-responsive genes across eight *Crocosphaera* genomes, including the six known *C*. *watsonii* strains and the coastal strains *C. subtropica* and *C. chwakensis*. In each genome, we identified all genes involved in P_i_ uptake, the hydrolysis of dissolved organic phosphorus (DOP), and the perception and regulation of gene expression in response to P_i_ limitation. Second, we analyzed the expression of these genes during P_i_ depletion in cultures of *C. watsonii* 8501. We also monitored the physiological parameters of cells in conditions of P_i_ depletion, as well as during a recovery phase in which DOP was provided to the P_i_-depleted cultures.

## Experimental procedures

### Strain and growth conditions

*Crocosphaera watsonii* strain WH8501, isolated in the western tropical South Atlantic Ocean, was grown under obligate diazotrophy in a YBCII culture medium (20), which we slightly modified to make sure the medium was devoid of any source of N (Fe-NH_4_-citrate was replaced with Fe-citrate). Cultures were kept at 27°C under a 12:12h dark:light cycle at 600 µmol photons.m^−2^.s^−1^ to ensure light-saturated growth. The growth medium was prepared from aged Mediterranean Sea water, collected at the Microbial Observatory Laboratoire Arago (MOLA) station (at 19 miles off the coast, 42°27.200’ N – 03°32.600’ E, 500m depth, salinity 38). The collected water was stored for at least 8 weeks in the dark at 20°C, filtered through 1 and 0.22µm Whatman filters, and autoclaved before use. Before the experiment, we analysed the bacterial contamination in the strain. Three strains of heterotrophic bacteria were identified: *Nitratireductor aquibiodomus*, *Qipengyuania goetbuli,* and *Oceaniradius stylonematis.* Bacterial contamination was then monitored during the experiments by flow cytometry with Sybr Green staining (see below). The bacterial contamination represented between 3.63 ± 1.23% and 7.7 ± 2.48% of the total biomass considering a carbon content of 4 or 8.9 fmol C.cell^−1^ as estimated in (21).

### Experimental design

A phosphate-replete culture (grown with 50 µmol L^−1^ of KH_2_PO_4_) was grown exponentially to serve as inoculum for all the experiment replicates. This culture, which reached an exponential growth rate of 0.26 d^−1^ (R^2^ = 0.95), was then filtered and transferred into fresh medium to prevent nutrient limitation and to allow a biomass build-up to the desired level. Once the biomass reached 10^7^ cells mL^−1^, the culture was filtered again and equally resuspended in 6 flasks containing fresh culture medium devoid of P_i_, aiming at an initial concentration around 3.5 10^6^ cells mL^−1^ in each flask. Due to the total estimated volume required for this experiment and the size limitations of the culture flasks, each replicate consisted of two flasks, each containing 2.5 L of P_i_-depleted culture. The six cultures were maintained under P_i_-depletion for a total of 9 days. All culture replicates were started simultaneously and monitored daily for cell abundance to verify that all population dynamics were equivalent. To ensure that the P_i_ remaining after dilution had been consumed and cells were experiencing P_i_ starvation, we waited for five days and initiated a high frequency monitoring from the 6^th^ day (labelled as Day 1 of monitoring on the figures) after the transfer to the P_i_-depleted medium. In total, cultures were monitored for 4 continuous days as follows: sampling started at the dark-light transition (L0) and was then performed every 3h, at L3, L6, L9, D0, D3, D6, D9, where L and D stand for light and dark, respectively. This sampling was performed in the first triplicate for the first 3 days; after the light-dark transition of the third day (Day 3, D0), the first replicate was nearly drained and sampling continued in the second triplicate.

Following the 9-day period in P_i_-depleted conditions, a recovery phase was initiated in the remaining (2^nd^) triplicate. The cultures were diluted in fresh, 2X (twice concentrated) YBCII medium devoid of phosphate and containing a mix of dissolved organic phosphorus (DOP), with a final concentration of 50 µmol P L^−1^ in the culture medium. We used an equal proportion of adenosine diphosphate, alpha glycerophosphate, nitrophenyl phosphate, and glucose 6 phosphate, each at a final concentration of 12.5 µmol P L^−1^ in the medium. To proceed with the DOP addition, 1 L of each of the remaining 3 replicates was sampled and transferred into an autoclaved, 3L erlenmeyer containing 750 mL of DOP enriched medium. After the DOP addition, the cultures were let to recover for 3 days and then monitored again at high frequency for 2 days.

### Analytical methods

#### • Cell size and abundance

*Crocosphaera* cell abundance was estimated using a Coulter Cytoflex (Beckman) flow cytometer by auto-fluorescent cell detection. Samples were taken for cell count at daily intervals before (during the P-replete growth phase) and after inoculation in a P_i_-depleted medium, and then every 3h during the high-frequency monitoring. The samples were fixed with glutaraldehyde (final concentration 0.5%) and conserved at −80°C until analysis. The contamination of the culture by heterotrophic bacteria was also verified by flow cytometry, in a second procedure after staining with SyberGreen. Thus, two acquisition procedures were applied based on the size and abundance differences between *Crocosphaera* and the heterotrophic bacteria. The growth rate was derived from cell counts by exponential regression of the abundance counts. The equivalent spherical diameter (ESD, µm) of each *Crocosphaera* cell was estimated by comparing the total forward scatter diffusion (the area under the FWS pulse shape) of the cells and that of manufactured Silica beads (known diameter of 1.05, 1.97, 3.93, 3.93µm; silica microsphere, Bangs Laboratories ®) every day of analysis. The particle-by-particle size data were extracted and analysed using FlowJo and R softwares. The biovolume (BV, µm^3^) was estimated with the following relation: 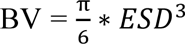 (Sun, 2003), with ESD the equivalent spherical diameter, and considering that *Crocosphaera* cells are spherical (5).

#### • Alkaline phosphatase activity

Alkaline phosphatase activity (APA) was monitored 3h after each light-dark transition: at L3 and D3, and was quantified using the fluorogenic substrate 4-Methylumbelliferone-Phosphate (MUF-P) (22) at a final concentration of 125 µmol L^−1^. Samples (210µL) were incubated in triplicate in black microplates at the culture temperature for 2 h. Control blanks were prepared from Milli-Q water with added MUF-P substrate. The fluorescence intensity was measured using a Perkin Elmer Victor3 Plate Reader fluorometer with an excitation wavelength of 365nm and an emission wavelength of 450 nm. Fluorescence readings were done at t_0_ (time of MUF-P addition), t_0_ +30 min, t_0_ +1 h, and t_0_ +2 h. The increases in fluorescence, confirmed to be linear during the incubations, were transformed into enzyme activity rates using a standard curve obtained from a MUF solution with concentrations ranging from 0 to 100 µmol.L^−1^.

#### • Particulate and dissolved nutrients

Samples for biochemical analyses were taken during the high-frequency monitoring phase. The particulate organic matter was determined every 6h, at L0, L6, D0, and D6. In each culture, two samples of 6.27mL were collected independently, one to analyze particulate organic carbon (POC) and nitrogen (PON) by complete combustion and the other for PON and particulate organic phosphorus (POP) quantification by wet oxidation. As PON was quantified with two distinct techniques, we differentiated PON_c_ obtained by combustion from PON_oxi_ obtained through wet oxidation analysis. Culture samples were filtered on pre-combusted (450°C - 4h) GF/C filters (nominal pore size 1.2 µm, Whatman). The filters were then dried and stored at 60°C before analysis. POC and PON_c_ filters were exposed to HCl fumes (4h) before analysis using a CHNS elemental Analyzer (Thermo Fisher Scientific) calibrated with acetanilide (23). PON_oxi_ and POP were estimated with a Segmented flux analyzer (Skalar), using the wet oxidation technique according to (24). C, N, and P cell contents (fmol cell^−1^) were deduced knowing the precise volume of the sample and the cell abundance. Because the two analytic techniques yielded different estimations of PON, we did not derive the biomass C:N:P stoichiometry. Instead, we expressed the C:N (deduced from the combustion analysis; mol:mol) and the N:P (deduced from the wet oxidation procedure; mol:mol) ratios separately.

Dissolved nutrients were sampled at L0 and D0 in each culture and conserved at −20°C before analysis. 30mL-samples were filtered on two GF/F combusted (450°C - 4h) filters (nominal pore size <0.5, Whatman). Filtrates of 7mL were used to quantify dissolved inorganic nitrogen (DIN) and phosphorus (DIP) according to Aminot et al. (2007) with a Segmented flux analyzer (Skalar). Standard solutions of NO_3_^−^ between 0 and 20 µmol.L^−1^ and blanks were prepared at the same salinity as the sample. Total dissolved nitrogen (TDN) and phosphorus (TDP) were analyzed with 10 mL of filtrate, according to the wet oxidation technique of Valderrama (1981), with a Segmented flux analyzer (Skalar). Duplicates of 500µL filtrate with 125 µL of reagent were used to quantify ammonium (NH_4_^+^). The fluorometric measure was realized 4 hours after sampling, according to (25) using a Denovix fluorometer with an excitation wavelength around 375nm and an emission wavelength between 435 and 485 nm. A calibration curve was constructed using concentrations ranging from 0 to 5 µmol L^−1^ in an NaCl solution at the same salinity as the sample.

### RNA sampling and preparation, reverse transcription, and quantitative Real-Time-PCR

Every 3h, 60 mL samples were collected and filtered through a 3µm pore-size polycarbonate membrane filter. The cells were resuspended in (0.2µm filtered and autoclaved) seawater, and centrifuged at 5000rpm for 15min. The pellet was transferred in 5mL RNA later, immediately frozen in liquid nitrogen, and stored at −80°C until analysis. All measurements were carried out in duplicate. RNAs were extracted using the Qiagen RNA-extraction kit following the manufacturer’s instructions. An extra TURBO DNase (Invitrogen) digestion step was performed to eliminate the contaminating DNA. The RNA quality was assessed by a tape station system (Agilent). RNAs were quantified spectrophotometrically at 260 nm (NanoDrop 1000; Thermo Fisher Scientific). For cDNA synthesis, 200 ng total RNA and 0.5 μg random primers (Promega) were used with the GoScript™ Reverse transcriptase (Promega) according to the manufacturer’s instructions. Quantitative real-time PCR (qPCR) analyses were performed on a CFX96 Real-Time System (Bio-Rad). The reaction volume was 15 μL and the final concentration of each primer was 0.5 μM. The qPCR cycling parameters were 98°C for 2 min, followed by 45 cycles of 98°C for 5 s, and 59°C for 10 s. The only exception was the 5’ND encoding gene, for which the melting temperature was 55°C. A final melting curve from 65°C to 95°C was added to determine the specificity of the amplification. To determine the amplification kinetics of each product, the fluorescence derived from the incorporation of SYBR^®^ Green Dye into the double-stranded PCR products was measured at the end of each cycle using the SsoAdvanced Universal SYBR^®^ Green Supermix 2X Kit (Bio-Rad, France). The results were analyzed using Bio-Rad CFX Maestro software, version 1.1 (Bio-Rad, France). The RNA 16S gene was used as a reference for normalization. The amplification efficiencies of each primer pair were 80 to 100%. All measurements were carried out in triplicate, a biological duplicate was performed for each point, and standard variations were calculated for each measure. The primers used in this study are listed in **Supporting information Table S1**.

### Bioinformatics analysis

#### Datasets

The genome data of 8 *Crocosphaera* strains available in September 2023 were downloaded from the NCBI ftp site (ftp:/ftp.ncbi.nih.gov/genomes/) and constituted the primary data source. This data source includes full genomes in the highest levels of assembly (complete genome, scaffold, or Contig), their RefSeq category (representative genome or none) as well as their annotation features (**Supporting information Table S2, Sheet 1**). Note that *Crocosphaera* species DT_26, ALOHA_ZT_9, and MO_202 were excluded from our set of genomes because these genomes are derived from metagenome assemblies. Cell-size features of *C. watsonii* (small and large) were provided by the literature (6). Hidden Markov Models (HMMs) of protein family profiles (Pfam version 35.0, April 2023) and Pgap (version 4, April 2023) were downloaded from ftp.ebi.ac.uk/pub/databases/Pfam/releases/ and ftp.ncbi.nlm.nih.gov/hmm/4.0/ ftp sites, respectively. Seed proteins (52 in total) used as references (retrieved from the Uniprot database, www.uniprot.org) in this study are described in **Supporting information Table S2, Sheet 2**. Proteins from cyanobacteria were used as probes when available; otherwise, proteins from *E. coli* were used as alternatives.

#### Functional domain search for orthologs identification

HMMER package (26) and HMM domain profiles were first used to establish the list of seed functional domains associated with reference seed proteins as well as their protein domain organizations (or domain patterns) (**Supporting information Table S2, Sheet 2**). Alignments with scores higher than the Pfam/Pgap trusted thresholds were considered significant seed domains. This led to 76 reference seed domains from Pfam or Pgap domain databases (**Supporting information Table S2, Sheet 3**). HMMER3 (26) and self-written Perl scripts were then used to search for protein orthologs (with seed domains) in the set of *Crocosphaera* genomes. For each reference protein, the presence of one or more seed domains was a requisite to identify putative orthologs. As described above, alignments with a score higher than the Pfam/Pgap thrust thresholds were also considered significant and in the case of overlapping domains, the longest ones with the best E-values were chosen. Putative orthologs of each seed protein were subsequently analyzed with the same software to determine their domain patterns with the potential presence of additional functional domains. Orthologs were then selected for the strict presence of seed domains (i.e., in the right order and without additional domain), and thus, led to the set of Hmm-homologs. To increase the sensitivity of the identification of orthologs, a Blast (27) analysis search was performed using all reference seed proteins as queries and an *E*-value threshold of 10^−5^. Blast alignments with a MinLrap [Alignment size / Minimum size (Query, Subject)] >= 0.8 and MaxLrap [Alignment size / Maximum size (Query, Subject)] >= 0.8 were considered as significant and constituted the set of Blast-orthologs. Hmm-homolog and Blast-homolog sets were finally combined to yield the final set of orthologs for each reference protein. For comparative analysis, reference proteins with identical Pfam domain patterns were considered identical seed proteins and then grouped as a unique reference for analysis. This was the case for PhnE, PstA, PstC, PtxC, UgpA, and UgpE [leading to PhnE-PstA-PstC-PtxC-UgpA-UgpE]; PhnC, PstB, PtxA, and PhnK [PhnC-PstB-PtxA-PhnK]; GppA and ppX [GppA-ppX]; PhnD and PtxB [PhnD-PtxB]; and PstS and SphX [PstS-SphX]. To further analyze the presence or absence of several key genes involved within the four systems known to be devoted to Pi limitation (Phosphate, Pst system; Phosphite, Ptx system; Phosphonate, Phn system; and Glycerol, Ugp system), tBlastN analyses were additionally performed. For that purpose, the corresponding seed proteins (associated to their localization: periplasmic, inner membrane or cytoplasmic) of each system were compared against our dataset of nucleotide genomic sequences with the E-value threshold was set to 10^−5^ for significant alignments and considering the best hit as the most likely ortholog.

## Results

### Functional genomics survey of P_i_-related genes in *Crocosphaera* genomes

To evaluate the genetic capabilities of various *Crocosphaera* strains in adapting to phosphate (P_i_) deficiency, a comprehensive search for homologs of proteins associated with transcriptional regulation in response to P_i_ deficiency, phosphate import, and dissolved organic phosphorus (DOP) hydrolysis was conducted. To achieve this, protein functional domain analysis, supplemented with BLAST search were performed for each category of proteins as described in the Experimental Procedures section. The findings for each functional category are outlined in the sections below and **Supporting information Table S2, Sheet 4**.

#### Signal transduction and gene regulation

Genes encoding orthologs to the histidine kinase sensor PhoR and the response regulator PhoB were detected in the genomes of the 8 analyzed *Crocosphaera* strains, suggesting that this two-component system (TCS) might, as in other bacteria, be the primary trigger of the P_i_ limitation response **(Figure 1, Supporting information Table S2, Sheet 4)**. The activity of the PhoR/PhoB system is controlled according to the concentration of phosphate in the environment. Under conditions where P_i_ is abundant, PhoU is essential for the dephosphorylation of PhoB (28) and plays a role in negatively regulating both the activity and the quantity of the Pst transporter to prevent excessive P_i_ uptake (29). This regulatory mechanism might be conserved in *Crocosphaera*, as an ortholog of the *phoU* gene was found in all the *Crocosphaera* genomes studied **(Figure 1, v)**.

**Figure 1.**
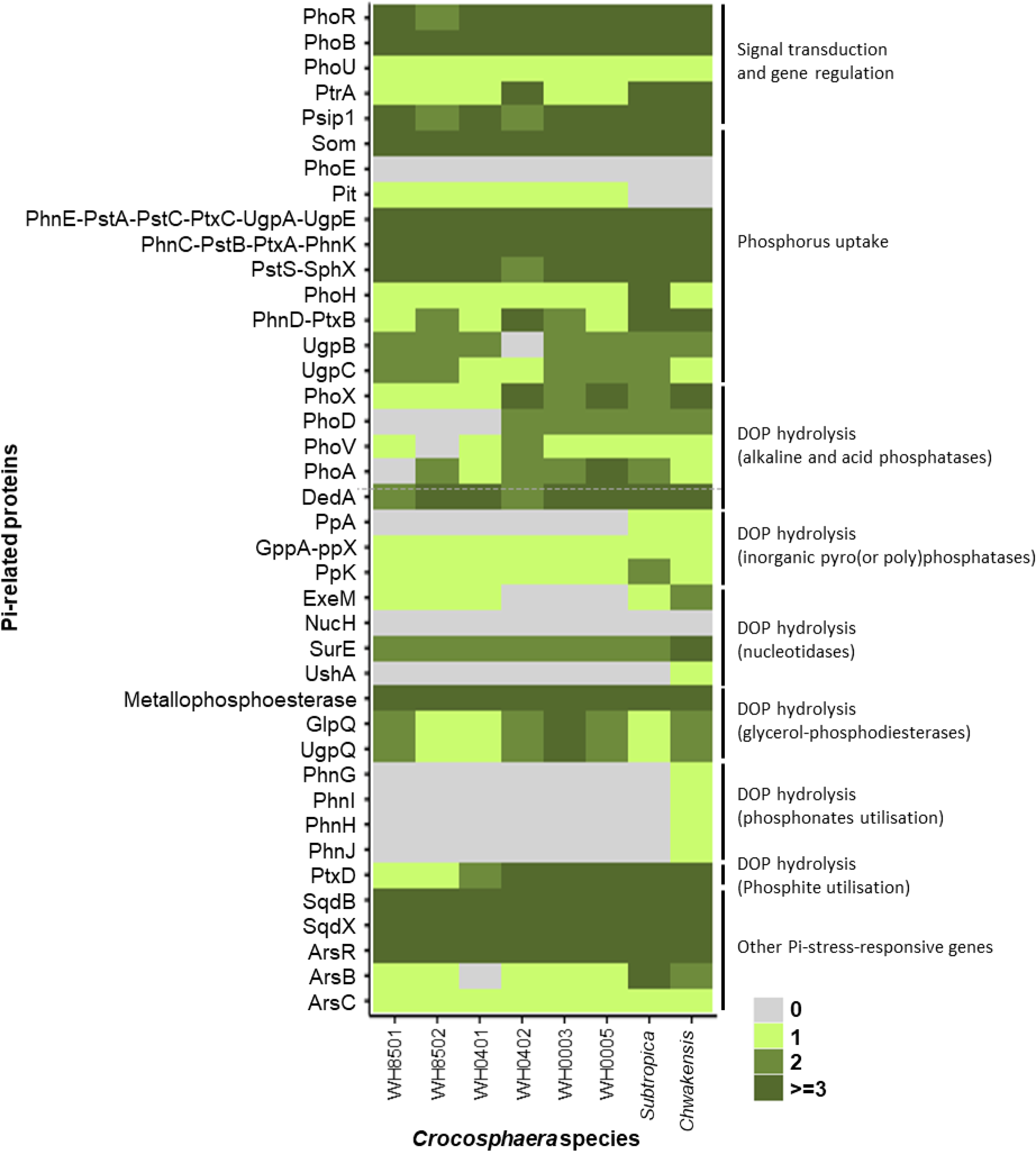
Heatmap representation of the number of P_i_-related gene orthologs within the 8 *Crocosphaera* genomes. The color scale and associated number from 0 to >= 3 represent the number of homologs found in each genome. Grey indicates that no ortholog sequence was found in the genome.

In the marine cyanobacterium *Synechococcus sp.* WH8102, the regulation of several P_i_-related genes involves a transcriptional regulator from the cAMP receptor protein (CRP) family, known as PtrA, that acts downstream from the PhoBR system (30). At least one copy of orthologs to *ptrA* was also found in all the *Crocosphaera* genomes **(Figure 1, Supporting information Table S2, Sheet 4).** Furthermore, a search for orthologs of the Psip1 protein from *Prochlorococcus marinus* WH8102, predicted to be part of the CRP family, revealed the presence of several copies of *psip1* genes in the *Crocosphaera* genomes **(Figure 1).** Altogether, these data indicate that CRP-like regulators probably have a significant role in coordinating the response to P_i_ stress in this genus.

#### Phosphorus uptake

Gram-negative bacteria can internalize P_i_ via non-specific porin channels in the outer membrane, facilitating the passive diffusion of small hydrophilic molecules. As an example, the major outer membrane protein Som in *Synechococcus sp. PCC 7942*, is involved in the uptake of these small molecules (31). In *Crocosphaera*, the P_i_ uptake through the outer membrane may also be achieved by the porin Som, as several copies of putative Som-encoding genes were found in all the *Crocosphaera* genomes studied (**Figure 1, Supporting information Table S2, Sheet 4**). Those *som* genes are located close to *phoU* and *phoR* genes, further suggesting a role of the Som porin in response to P_i_ stress. In *E. coli*, the porin PhoE, which is part of the Pho regulon, facilitates the uptake of P_i_ across the outer membrane (32). No PhoE ortholog was identified in the protein database of the *Crocosphaera* strains studied (**Figure 1, Supporting information Table S2, Sheet 4**), suggesting that a *phoE* gene is unlikely to exist in this cyanobacterial genus.

To identify proteins that may facilitate P_i_ uptake across the cell inner membrane, we searched for orthologs of the Pit system, which facilitates P_i_ entry under non-limiting conditions, and the Pst system, which is activated under low P_i_ concentrations (15). Orthologs of the Pit protein were found in the protein database of *C. watsonii* strains, but intriguingly, not in *C*. *subtropica* and *C. chwakenis*. The absence of Pit orthologs is puzzling; given their coastal habitats, these strains are unlikely to consistently face phosphorus limitation so we expected to find orthologs of the low-affinity transport system Pit. Whether they utilize an alternative system for constitutive P_i_ uptake therefore remains to be investigated. The Pst system is an ATP-binding cassette (ABC) transporter comprising three essential units: PstC, PstA, and PstB. These are encoded by the *pstSCAB* gene operon, which belongs to the Pho regulon. PstS, a periplasmic P_i_ binding protein, delivers P_i_ to the transmembrane proteins PstC and PstA. PstB, a cytosolic ATPase, uses ATP hydrolysis to import P_i_ into the cytoplasm (10). In cyanobacteria, a PstS homolog, designated SphX, belongs to the phosphate-binding protein family and delivers periplasmic P_i_ to the ABC transporter (33).

In addition to P_i_, phosphites, phosphonates, and glycerol-phosphate can be used as sources of phosphorus by bacteria. The uptake of these molecules through the inner membrane is achieved by ABC transporters which belong to the same family as the Pst system. Phosphite is incorporated through the Ptx system, phosphonate through the Phn system, and glycerol-phosphate through the Ugp system. Each subunit of a given transporter shares the same functional domain signature with its orthologs in the other systems, except for UgpC and UgpB, which have distinctive functional domains (**Supporting information Table S2, Sheet 2)**. Consequently, using functional domains as a search tool to identify orthologs does not allow the identification of subunits specific to each transporter. This analysis yields all potential orthologs for each subunit group, as shown in the heatmap of **Figure 1, Supporting information Table S2, Sheet 4**.

To specifically predict the presence of a complete set of genes for each transporter, we completed our analysis using tBlastN. We considered the ortholog with the best hit for each search as the most probable ortholog. The results indicate that all eight *Crocosphaera* possess a complete set of genes for the Pst system **(Figure 2 and Supporting information Table S2, Sheet 5).** It should be noted, however, that in the case of *C. watsonii* WH8501, the *pstA* gene is considered a pseudogene in NCBI database, which results in the absence of the corresponding protein in the Refseq database. Therefore, the *pstA* gene cannot be retrieved from WH 8501 genome. The annotated pseudogene may be due to sequencing errors which may have interrupted the open reading frame of this gene. Alongside the Pst system, the gene *phoH*, which encodes a putative ATPase, is found in the Pho regulon of bacteria, although its exact function remains unclear (34). Orthologs of PhoH were found in all *Crocosphaera* genomes, suggesting that this protein may play a role in adaptation to Pi limitation **(Figure 1**).

**Figure 2.**
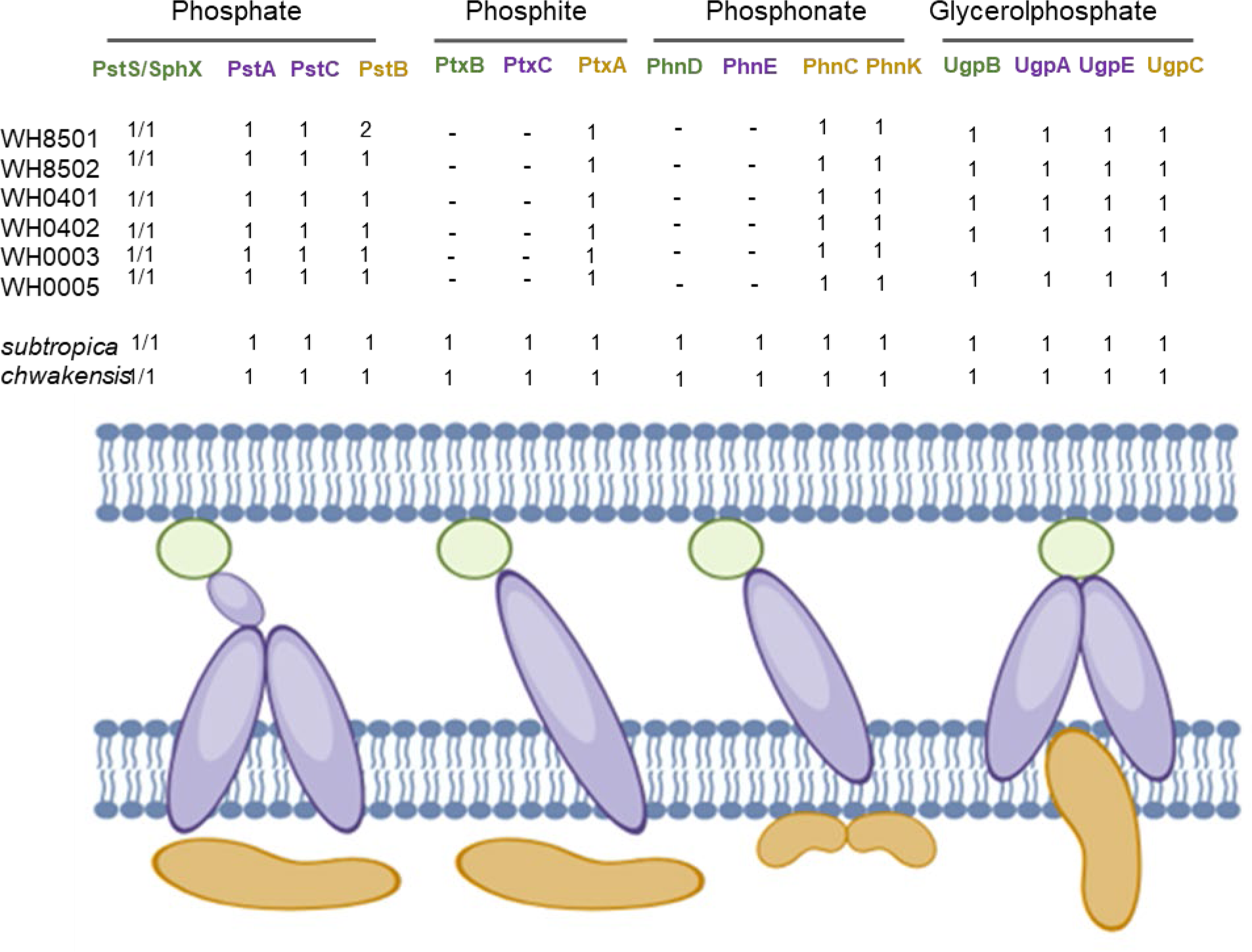
Schematic representation of the distribution of P_i_ transporters in *Crocosphaera.* 1 and 2 indicate the number of orthologs in each genome. (-) stands for the absence of ortholog. In each system, the P_i_ binding periplasmic protein is in green, the membrane subunit(s) in purple, the cytoplasmic ATPase subunits in brown. The figure was drawn using BioRender

Gene orthologs encoding all the subunits of an Ugp transport system were identified in all the *Crocosphaera* genomes, suggesting that all eight strains may be capable of internalizing phosphoglycerol **(Figure 2 and Supporting information Table S2, Sheet 5).** In contrast to the Pst and Ugp systems, only the genomes of *C. subtropica* and *C. chwakensis* possess a complete set of genes encoding all the subunits of the Ptx and Phn transporters **(Figure 2 and Supporting information Table S2, Sheet 5)**. Taken together, these genome analyses suggest that all *Crocosphaera* strains may be able to use phosphate and glycerol-phosphate. A discrepancy is found within the genus regarding the use of phosphites and phosphonates, with the six *C. watsonii* strains being likely unable to uptake them.

#### DOP hydrolysis

We carried out an extensive analysis in search of a wide variety of genes known to encode enzymes capable of generating phosphate from various substrates, such as phosphoesters (alkaline and acid phosphatases), polyphosphates (polyphosphatases), pyrophosphate inorganic pyrophosphatase), and nucleotides (nucleotidases) (**Supporting information Table S2, Sheet 2).** Of all the alkaline phosphatases known in bacteria, only the *phoX* gene has been identified in all the *Crocosphaera* genomes. No ortholog to the *phoD* gene was found in the *C. watsonii* WH8501, WH8502, and WH0401 genomes. The *phoV* gene was not detected in the *C. watsonii* WH8502 genome and no ortholog to *phoA* was identified in *C. watsonii* WH8501. Consequently, all *Crocosphaera* strains are potentially able to hydrolyze DOP since their genomes contain at least one gene encoding a typical alkaline phosphatase (**Figure 1, Supporting information Table S2, Sheet 4**.). In addition, all their genomes contain an ortholog of the *dedA* gene encoding an alkaline phosphatase-like enzyme.

The use of inorganic pyrophosphate as a source of phosphate is probably limited to the *C. subtropica* and *C. chwakensis* strains since an ortholog to the *ppA* gene encoding a pyrophosphatase was only found in their respective genomes. Conversely, all strains might be able to hydrolyze polyphosphates since at least one ortholog encoding a polyphosphatase (PpX or GppA) was found in their genome. In addition, the eight *Crocosphaera* strains might synthesize polyphosphate reserves, as suggested by the presence of the *ppK* gene potentially encoding a polyphosphate polymerase catalyzing polyphosphate formation from ATP.

Three nucleoases (NucH, SurE, and UshA) were identified in our study. While a SurE ortholog was identified in all the genomes analyzed, only one UshA ortholog was identified in *C. chwakensis* and the NucH gene was absent in all 8 genomes (**Figure 1, Supporting information Table S2, Sheet 4**.). The SurE 5’ nucleotidase could be used for phosphate recycling in the *Crocosphaera* genus. It is worth noting that this gene is annotated as *phoA* in some *Crocosphaera* genomes, and this annotation error may mislead the interpretation of functional measurements of phosphatase alkaline activities (see the Discussion section). Our results also indicate that the genus *Crocosphaera* could use phosphoglycerol as a source of phosphate since orthologs to the *glpQ* and *ugpQ* genes encoding glycerol-phosphodiesterases are present in the 8 genomes analysed. The ability to use phosphonates is likely restricted to *C. chwakensis* as a complete set of genes for hydrolysis of these compounds (*phnGHIJM*) was only found in its genome.

Interestingly, although only one strain of *Crocosphaera* seems to have a phosphite entry system (as mentioned above), the *ptxD* gene responsible for phosphite utilization was identified in all 8 genomes analyzed. This point raises an intriguing question: did the evolution of phosphite use occur after the loss of genes encoding transporters in most strains, or are these genomes currently acquiring these genes?

#### Other P_i_-stress-responsive genes

Substituting phospholipids with sulfolipids is a strategy cyanobacteria may use to cope with P_i_ scarcity (35). The presence of orthologs to the s*qdB* and *sqdX* genes, which encode two essential synthases required for glyceroglycolipid synthesis, in all *Crocosphaera* genomes (**Figure 1, Supporting information Table S2, Sheet 4**.) suggests that this adaptive strategy might be conserved within this genus.

Arsenate is a toxic substance coincidentally internalized by the P_i_ transporters due to its similarity to P_i_. As a result, inducing arsenate-reduction activities is part of the adaptive responses to P_i_ limitation (36). The eight *Crocosphaera* strains appear to be able to detect the presence of arsenate, as their genomes contain an ortholog of the *arsR* gene, which encodes the specific transcriptional regulator for genes encoding the arsenate reductase ArsC and the efflux pump ArsB. Although the *arsC* gene is present in all *Crocosphaera* genomes, *arsB* is absent in the *C. watsonii* WH0401 (**Figure 1, Supporting information Table S2, Sheet 4**.). Arsenate reduction might be the unique strategy to cope with its toxicity in this strain.

The relationship between morphotype and gene conservation was examined using three groups: the *C. watsonii* small-cell strains (WH0401, WH8501, and WH8502), the *C. watsonii* large-cell strains (WH0003, WH0005, and WH0402), and the ancient *Cyanothece* spp., newly classified within the *Crocosphaera* genus as *C. subtropica* and *C. chwakensis*. The gene encoding the low-affinity P_i_ transporter, *pit*, was identified as an ortholog in the *C. watsonii* but not in the new *Crocosphaera* strains. Conversely, the new *Crocosphaera* strains were the only ones to possess genes encoding the phosphonate transporter (*phnGHIJ*), inorganic pyrophosphatase (*ppA*), and 5’-nucleotidase (*ushA*) (**Figure 3, Supporting information Table S2, Sheet 5**,**6**). Overall, the percentage of total orthologs was lowest in the small-cell strains compared to the other two groups. For example, the small-cell strains lacked orthologs for the potential alkaline phosphatase (AP) gene *phoD*, while more sequences encoding AP (*phoA, phoD, phoX, and pafA/phoV*) were found in the large-cell and new *Crocosphaera* strains (**Figure 3, Supporting information Table S2, Sheet 5**,**6)**.

**Figure 3.**
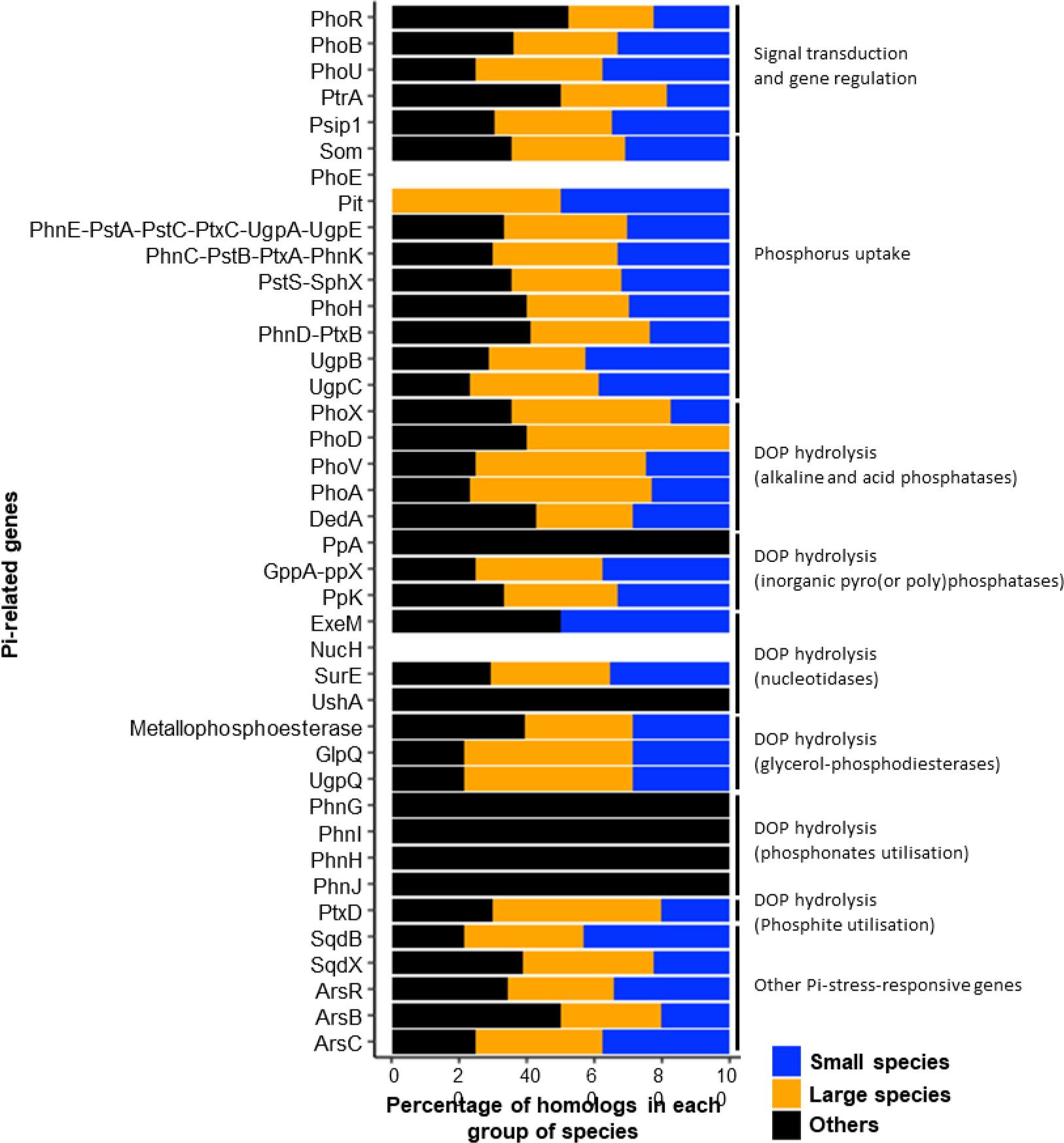
Percentage of P_i_-related gene orthologs in group of species. The color code based on cell size is shown. Small-cell strains (blue; *C. watsonii* WH8501, WH8502 and WH0401), large-cell strains (orange; *C. watsonii* WH0402, WH0003 and WH0005) and the newly classified strains in the genus (black; *C. subtropica* and *C. chwakensis)*.

### Diel expression of genes involved in P_i_-depleted and DOP-recovery conditions

To investigate the genetic response of *Crocosphaera* to phosphorus-depleted (P_i_-depleted) and DOP recovery conditions, we analyzed the transcription of P_i_-related genes every 3 hours over 4 days under P_i_-depletion and then over 2 more days after the same cultures were supplied with DOP.

The expression of *ptrA*, which encodes a potential regulator in response to P_i_ limitation, was low during the Pi depletion phase and followed a daily cycle with peaks during the light phase. Specifically, the peaks occurred at L9 on Day 2 and L6 on Day 4 (**Figure 4A**). 3 days after DOP addition, *ptrA* expression was three times lower than the first sampling point (L0, Day 1) of the P_i_-depleted phase (**Figure 4A**).

**Figure 4.**
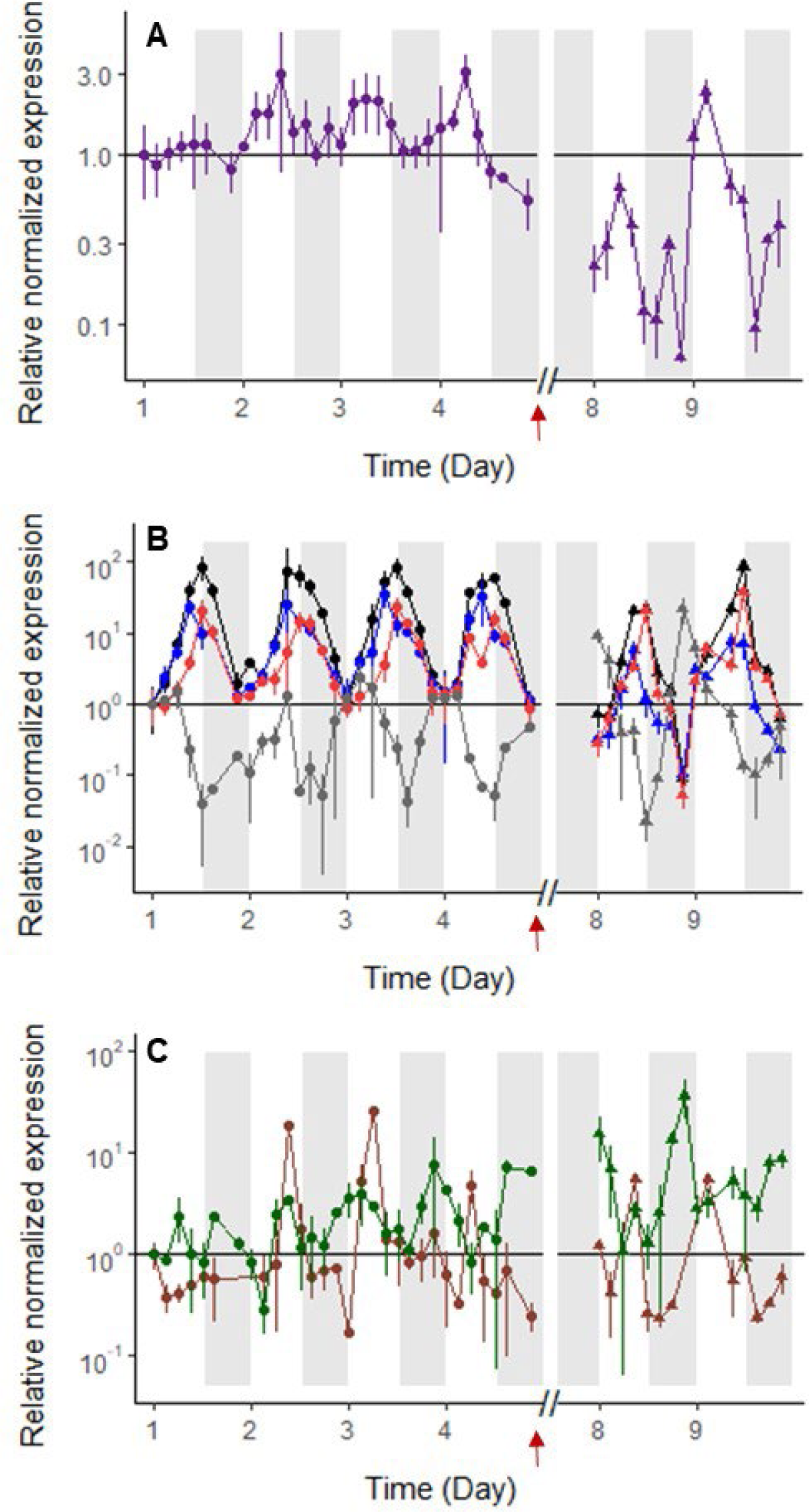
Relative normalized gene expression of *C. watsonii* WH8501 during the P_i_-depleted phase (closed circles, days 1-4) and the DOP-recovery phase (closed triangles, days 8-9). RT-qPCR of the genes encoding for a regulator *ptrA* in purple (**A**), for a porin *som* in red, and transporters, *sphX* in black, *pstS* in blue, and *ugpC* in grey (**B**). Expression of the genes encoding for a glycerophosphoesterase *ugpQ* and a 5’-nucleotidase are represented in brown and green, respectively (**C**). The Y-axis scale is logarithmic. Time on the X axis is expressed in days, starting from the beginning of the high-frequency monitoring phase, i.e. five days after the transfer to a P_i_-depleted medium. All points are normalized by the 16S expression at the same point and relative to the first sampling point (Day 1, L0). The horizontal line (y=1) materializes the value of the first sampling point. The expression variability of biological and analytical duplicates is represented by error bars. White and grey shades represent light and dark periods, respectively and red arrows mark the time of DOP addition.

Given that genes encoding P_i_ transporters are organized in clusters, suggesting a possible operonic organization, we chose to analyze the expression of the first gene, encoding the binding protein, in each cluster as a representative of the entire gene cluster. The genes encoding the binding protein of the high-affinity P_i_ transporter (PstS) followed a daily expression cycle, peaking at the light-dark transition (D0). The *pstS*-homolog *sphX* gene peaked three hours before the light-dark transition (L9) in the P_i_-depleted phase **(Figure 4B**). The gene encoding the glycerol-phosphate molecule binding protein (UgpC) showed a low but cyclic expression, with a higher expression during the light phase than the dark phase (**Figure 4B**). The porin Som-encoding gene displayed the same expression profile as *pstS*. During the DOP-recovery phase, the expression peaks of *som*, *pstS*, and *sphX* occurred at the same times as during the P_i_-depleted phase. However, the expressions of *pstS* and *sphX* were lower compared to the P_i_-depleted phase (four times lower on Day 8 for *sphX*, and on Days 8 and 9 for *pstS*), whereas *ugpC* expression was ten times higher on Days 8 and 9 compared to Day 1 (**Figure 4B**).

Genes potentially encoding enzymes required for P_i_ scavenging were expressed globally at a low level (**Figure 4C and Supporting information Figure S1**). During the P_i_-depleted phase, the genes encoding the glycerophosphoesterase UgpQ and the 5’-nucleotidase (5’ND) showed oscillatory patterns (**Figure 4C**). UgpQ transcription peaks in the light phase were over ten times higher than the initial point on Days 2 and 3 (**Figure 4C).** The expression of the 5’ND encoding gene peaked during both light and dark phases and increased over tenfold during the DOP-recovery phase (**Figure 4C**). Other genes encoding enzymes, such as the potential 5’nucleotidase SurE, a metallophosphoesterase, the alkaline phosphatase-like (DedA) (**Supporting information Figure S1**), and the alkaline phosphatase PhoX, were lowly expressed throughout the experiment (data not shown). In conclusion, the expression of genes potentially encoding a high-affinity P_i_ transporter exhibited diel variations during both the P_i_ depletion and DOP recovery phases. In contrast, genes predicted to encode DOP-hydrolyzing enzymes were expressed at low levels and displayed different patterns between the two growth phases.

### Expression of circadian clock genes

The observed daily cyclical gene expression may be driven by circadian control exerted by an internal clock. Initially identified in the unicellular cyanobacterium *Synechococcus elongatus* PCC7942 (37), the circadian clock is largely conserved within cyanobacteria (38). But, to our knowledge, the circadian clock has not yet been studied in *C. watsonii* WH8501. To investigate whether the cyclic expression of the genes studied is linked to a circadian control, we searched for homologs of the *kaiA*, *kaiB*, and *kaiC* genes, which constitute the *Synechococcus* clock, as well as the *rpaA* gene, which encodes a transcriptional regulator that transduces the control generated by the clock (39).

Homologs of the *kaiABC* genes were identified in the *C. watsonii* WH8501 genome, clustered together (**Figure 5A**). In *Synechococcus*, the *kaiB* and *kaiC* genes form an operon, and all three *kai* genes exhibit circadian expression (39). In the *C. watsonii* WH8501 genome, the *kaiA* and *kaiB* genes are contiguous, suggesting they belong to the same transcriptional unit. Analysis of the transcription of these three genes under the experimental conditions described above reveals an intriguing profile. While the *kaiB* and *kaiC* genes showed circadian expression with a transcriptional peak at light-dark transition, the expression of the *kaiA* gene did not appear to be circadian and followed a different profile from that of *kaiB* (**Figure 5B, C, D)**. These findings support the hypothesis that the clock controls gene expression in *C. watsonii* WH8501. However, they also suggest that the functioning of this clock may differ from that in *Synechococcus*, calling for further investigation. Additionally, the transcription of the *rpaA* gene was cyclic, with peak activity during the night phase (**Figure 5D**), consistent with its predicted role in the output system acting downstream of the clock.

**Figure 5.**
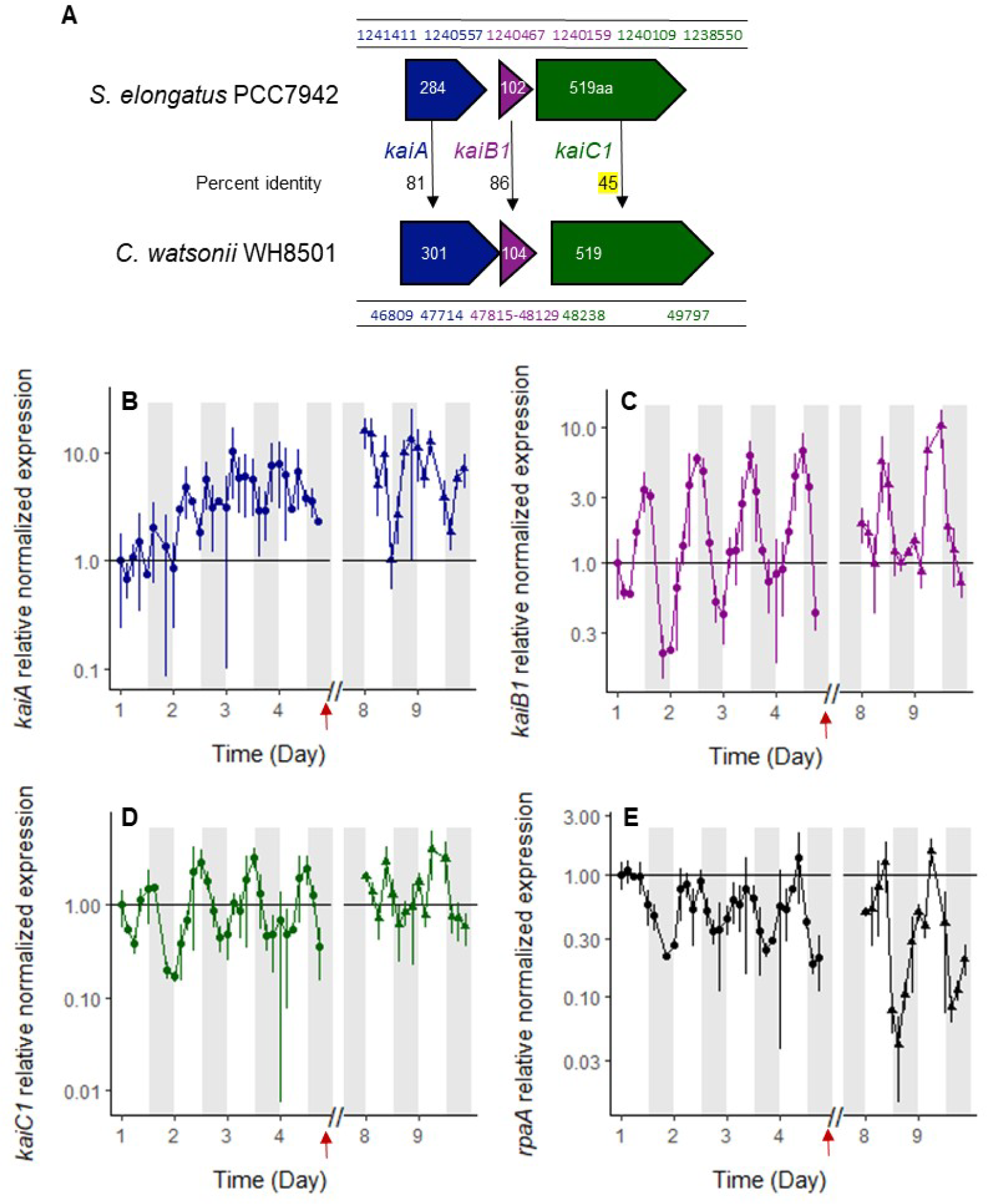
Gene cluster organization and genomic position of *Crocosphaera watsonii* WH8501 potential *kaiABC* and percent identity to *Synechococcus elongatus* PCC 7942 (**A**); relative normalized expression of *C. watsonii* potential circadian clock components *kaiA* (**B**), *kaiB* (**C**), *kaiC* (**D**) and regulator, *rpaA* (**E**) during the P_i_-depleted phase (closed circles) and the DOP-recovery phase (closed triangles). The Y-axes are logarithmic. Time on the X axis is expressed in days, starting from the beginning of the high frequency monitoring phase, i.e. five days after the transfer to a P_i_-depleted medium. All points are normalized by the 16S gene expression at the same point and relative to the first sampling point (Day 1, L0). The horizontal line (y=1) materializes the value of the first sampling point. Expression variability of biological duplicates and analytical duplicates are represented by error bars. White and grey shades represent light and dark periods, respectively, and red arrows mark the time of DOP addition.

### Population dynamics

We followed the cell abundance during P starvation and DOP recovery by flow cytometry to compare the changes in population dynamics and quantify the growth rates. Upon transfer of the exponentially growing cultures to a P_i_-depleted medium, the initial cell concentration in the P_i_-depleted replicates was 3.49 ± 0.22 10^6^ cells mL^−1^ (n=6). A transient increase in cell abundance was visible in all cultures, which grew at a rate of 0.23 ± 0.03 d^−1^ on average (n=6 replicates with 5 time-points) over the first four days (days −4 to 0, **Supporting information Figure S2**). This rate is slightly lower than the rate observed in the seed culture grown on P_i_ (0.26 d^−1^) but it was also estimated in denser cultures, which most probably experienced a slightly lower irradiance compared to the seed culture. In that respect, the initial growth rate in the P_i_-depleted phase was comparable to that observed in the seed culture grown with P_i_. In contrast, the exponential phase appeared much shorter in time. An inflection in the temporal fluctuation of the cell abundance was visible from the 5^th^ day into the P_i_ depleted phase (day 1); the biomass kept increasing for another couple of days, albeit at a slower pace, and reached a plateau with an average cell abundance of 1.05 10^7^ cells mL^−1^ in the 6 replicates (**Supporting information Figures S2 and 6A**). Overall, in 8 days, the cell number tripled and cells divided 1.6 times on average, resulting in an average growth rate of 0.14 d^−1^ over the 8 first days of P_i_ depletion. Cell abundance thereafter initiated a decrease from the 9^th^ day into the P_i_-depleted phase (day 4) when the DOP-containing medium was added to the 3 remaining replicates. Cell counts recorded on the 4^th^ and 5^th^ day following the DOP addition indicate that cells started growing again (**Figure 6A**), at a rate of µ_+DOP_ = 0.31 ± 0.01 day^−1^ (n=16 for each replicate; average from the three estimated rates) over the two days of monitoring. We do not consider this rate very accurate as it derives from two consecutive days only and the doubling time at this growth rate is more than two days. However, the consistency of the estimates between the three replicates as well as the clear difference in population dynamics compared to the P_i_-depleted phase are solid proof for the good recovery of the strain and the use of (some, at least) of the DOP compounds provided to re-initiate growth.

**Figure 6.**
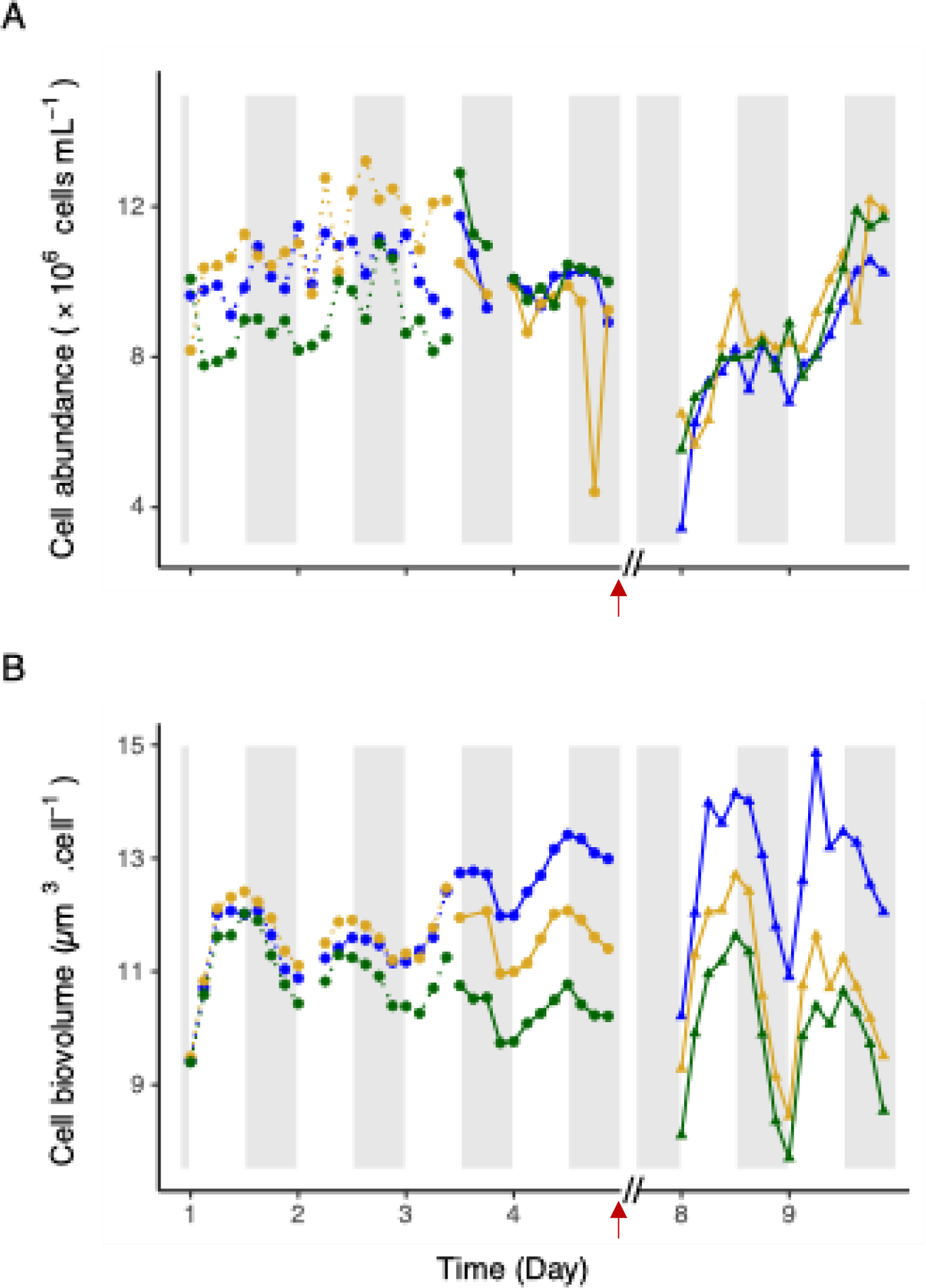
Diel fluctuations in *C. watsonii* cell abundance (top panel, 10^6^ cells mL^−1^) and cell biovolume (bottom panel, µm^3^ cell ^−1^) represented for each culture replicate (blue, yellow, and C: green). Time on the X axis is expressed in days, starting from the beginning of the high-frequency monitoring phase, i.e. five days after the transfer to a P_i_-depleted medium. The dotted lines represent sampling in the first culture triplicate and continuous lines represent samples taken in the second triplicate (see methods). The red arrow marks the time of DOP addition in the second triplicate; sampling of the DOP-recovery phase was performed during Days 8 and 9. White and grey shades represent light and dark periods, respectively.

Under non-limiting nutrient conditions, cell size is known to follow a diel cycle of increase between the end of the dark phase and the end of the light phase, and decrease during the dark phase (40) (41). Under the present, transient conditions of P_i_ starvation, cell size kept oscillating with a similar tendency to increase in the light and decrease in the dark. The timing of these oscillations was very consistent between all replicates, albeit with diverging average sizes during the P_i_ depletion. The average cell diameter was 2.79 ± 0.07 µm (n=92). Without sufficient size records under the initial, phosphate-replete growth, we cannot assess whether the P_i_ depletion affected the average cell size. The measured values still fall within the ranges usually observed for this strain (42) (41) (43). The corresponding biovolume was 11.94 ± 0.89, 11.60 ± 0.60, and 10.70 ± 0.62 μm^3^ cell^−1^, with an overall average of 11.41 ± 0.88 µm^3^ cell^−1^ (n=92) (**Figure 6B, days 1 to 4**). On the fourth day following the refreshment of cultures into the DOP-containing medium (days 8 and 9), the amplitude of diel oscillation in cell size had doubled. The average cell diameter was 2.90 ± 0.10 (n=16), 2.74 ± 0.11 (n=16), and 2.66 ± 0.11 μm (n=16) in the three replicates, with an overall average of 2.77 ± 0.14 µm (n=48). The corresponding biovolume was then 12.86 ± 1.24, 10.80 ± 1.25, and 9.91 ± 1.19 μm^3^ cell^−1^ in the three replicates, with an overall average of 11.19 ± 1.73 µm^3^ cell^−1^. This much wider fluctuation in cell size at the diel scale suggests a recovery of carbon fixation, accumulation in the light, and consumption in the dark.

In addition to the diel oscillation, a transient decrease in cell size also normally occurs around the mid-light phase in nutrient-replete cultures, consequent to cell division (40, 41). In the present experiment, no decrease in cell diameter can be observed around mid-light during the P_i_-depleted phase, while it becomes visible during the DOP recovery. This is another evidence for the disruption of cell division towards the end of the P_i_-depleted phase and for the recovery of growth and cell division after cultures were provided with DOP. (**Supporting information Figure S2**)

### Dissolved macronutrients and alkaline phosphatase activity

We monitored dissolved N and P concentrations in the cultures and DOP hydrolytic activities to track the consumption of nutrients, which also informs on growth efficiency. Nitrate (NO_3_^−^) concentrations were close to zero (0.36 ± 0.28 µmol N L^−1^) during the whole experiment (**Table 1**) since the medium did not contain any added nitrate. Organic (DON) and total dissolved nitrogen (TDN) levels were both higher during the DOP-recovery phase compared to the P_i_-depleted phase (19.10 ± 1.70 *vs* 9.19 ± 0.91 µmol N L^−1^ for DON and 19.68 ± 1.74 *vs* 9.44 ± 0.91 µmol L^−1^ for TDN; **Table 1**). Most likely, N exudation processes were strongly reduced under P_i_ depletion and restored during the recovery phase. The exuded DON in the recovery phase may also be constituents of DOP hydrolytic enzymes.

**Table 1.**
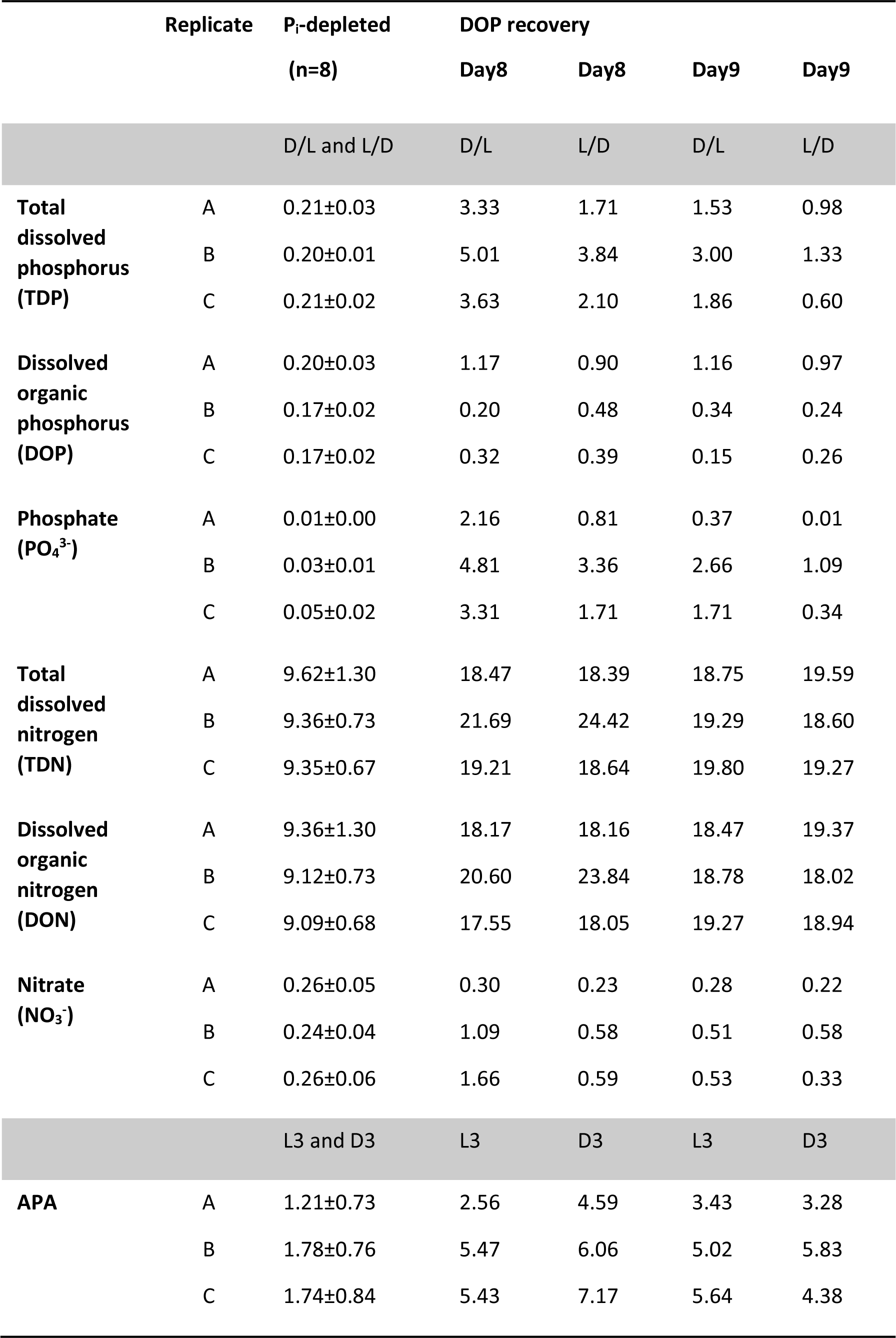
Dissolved nutrient concentrations (µmol N or P L^−1^) and per-cell alkaline phosphatase activities (APA, fmol cell^−1^ h^−1^) in replicates A, B, and C. In the P_i_-depleted phase, values varied little between dark-light (D/L) and light-dark (L/D) transitions, and so were averaged ± standard deviation in the culture over the entire monitored period. In the DOP-recovery phase, macronutrient concentrations are reported for the dark-light (D/L) and light-dark (L/D) transitions of days 8 and 9, and the APA at L3 and D3.

During the P_i_-depleted phase, the measured total dissolved P (TDP) in all cultures showed mean concentrations as low as 0.21 ± 0.02 µmol P L^−1^, with 0.03 ± 0.02 µmol P_i_ L^−1^ and 0.18 ± 0.03 µmol DOP L^−1^ (**Table 1**). Three days after the DOP addition (days 8 and 9), DOP concentrations had dropped to a range between 0.15 and 1.17 μmol P L^−1^ while phosphate levels were between 2.16 and 4.81 μmol P L^−1^ on day 8 and showed a decreasing trend over the two days monitored (**Table 1**). These results suggest that most of the DOP added was rapidly hydrolyzed and the liberated Pi was mostly incorporated in cells. The alkaline phosphatase activity assay (APA) was used to assess the production and activity of DOP-hydrolyzing enzymes. The expression of alkaline phosphatases is part of the cell response to phosphate limitation in cyanobacteria (44) (45) (46) (47), including diazotrophic strains (15) (14). The APA yielded rather low and stable activities during the P_i_-depleted phase, with an overall average recorded activity of 1.56 ± 0.79 fmol cell^−1^ h^−1^ (n = 22). Rates increased 3.15-fold during the DOP-recovery phase, with an overall average of 4.90 ± 1.32 fmol cell^−1^ h^−1^ (n = 12) (**Table 1).**

### Particulate carbon, nitrogen, phosphorus, and C:N:P ratios

The total *Crocosphaera* C, N, P biomass and its stoichiometry were followed and normalized by the population abundance to derive cellular contents. We consider all particulate C, N and P to be part of the cells. Exuded EPS may also be collected on filters but we consider the contribution of EPS to be very small in these thin, monocultures. During the four days monitored in the P_i_-depleted phase, the cellular carbon (POC) and nitrogen (PON) contents fluctuated somewhat irregularly, without any evident diel pattern reproduced from one day to the next (**Figure 7A, B, C**). The last record in replicate B, which stands as an outlier in this data set, is not considered in the following. Although irregular, the variations in the carbon cell contents remained in a rather conserved interval between 259 (163 if considering the first point that also stands out of the general trend) and 434 fmol C cell^−1^, with a very slight decreasing trend of 8 fmol C lost per cell and day on average (**Figure 7A**). The cellular nitrogen content (PON_c_) also varied in a rather conserved range between 26 and 45 fmol N cell^−1^, with a decreasing trend in time of 1.89 fmol N lost per cell and day (**Figure 7C**). Carbon and nitrogen acquisition occurred but could not completely compensate for the losses. Compared to the steady, diel oscillations in the C and N contents reported in exponentially growing cultures (48) (40) (41) (49), the more erratic, transient changes in the present data suggest that both the C and N metabolism were partly impaired over those days. The resulting C:N ratio still showed the diel oscillations known for this strain that result from the temporal decoupling of C and N acquisitions. However, the increasing trend drifting far above the Redfield (1934) (50) canonical reference (with values between 10 and 12 on the last day of the P_i_-depleted phase) reveals a skewed N stoichiometry: N incorporation was proportionally more impaired than C fixation. Data also suggest that cells in one culture replicate tended to present higher C, N and P contents than the other two (**Figure 7A-D**, blue curve). But cells were also somewhat bigger in that replicate (**Figure 6B**); as a result, C and N content normalized by the biovolume showed more similar values in all three replicates (**Supporting information Figure S3**). The cellular P content fluctuated between 0.72 and 1.69 fmol P cell^−1^, considering the last point of replicate B as an outlier. The null derivative of this trend suggests that the average P content in cells remained rather constant over the four days monitored in the P_i_-depleted phase (**Figure 7D**). The resulting mol:mol N:P ratio varied with no obvious diel trend, between 17.0 and 28.5, and showed an average value of 22.9 ± 2.9 (n = 47) (**Figure 7F**). This average is markedly above the Redfield reference, indicating that the cellular P content is particularly low with respect to N. Altogether, these results suggest that under severe phosphate depletion, the P stoichiometry strongly drops and the N stoichiometry is also reduced, while carbon acquisition seems proportionately less affected.

**Figure 7.**
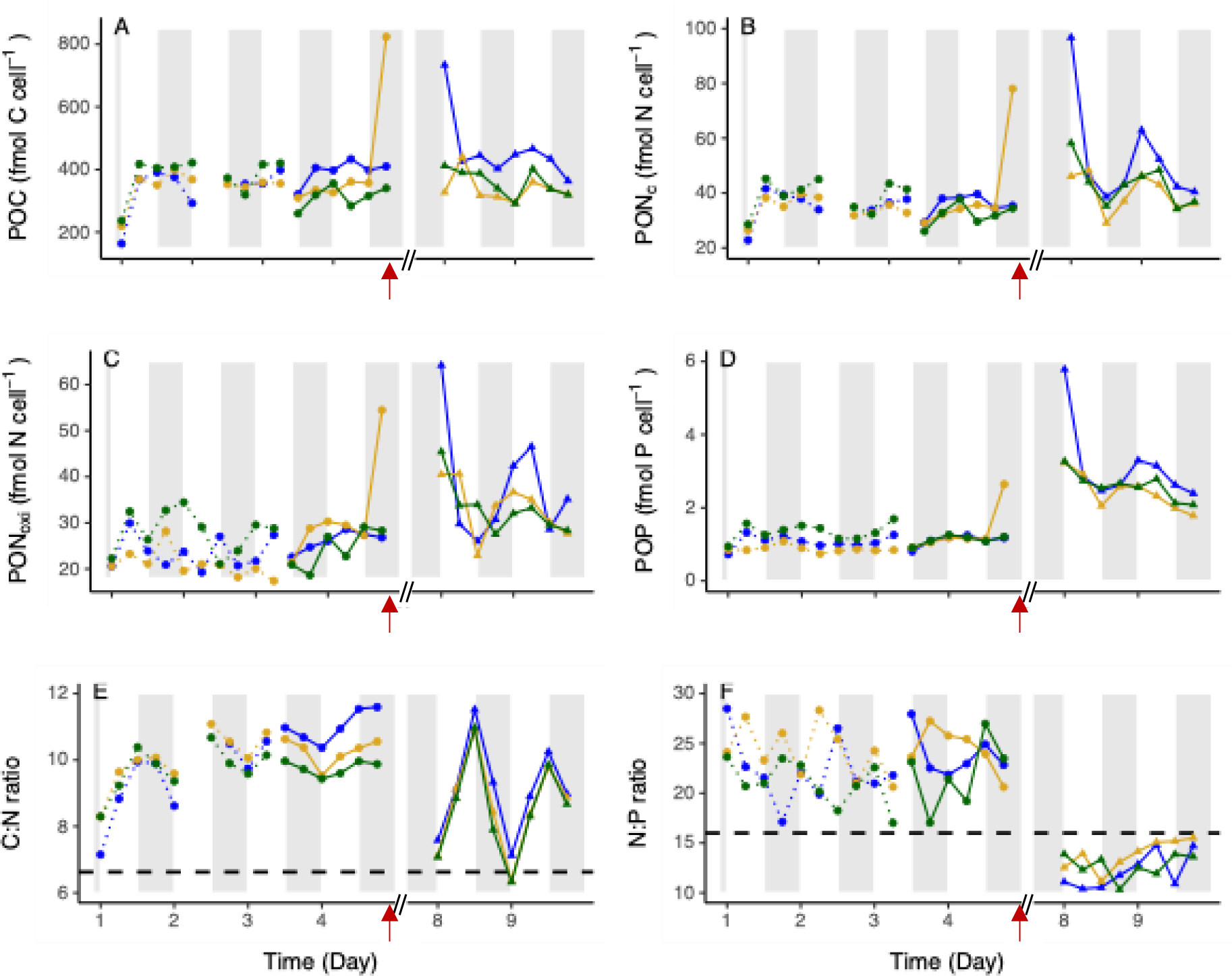
Diel fluctuations of *C. watsonii* C, N and P cell contents in each replicate (blue, yellow and green) during the P_i_-depleted (Day 1 to Day 4) and DOP-recovery (Day 8 and Day 9) phases. Particulate organic carbon (POC, fmol C cell^−1^, A) and particulate organic nitrogen (PON_c_, fmol N cell^−1^, C) were used to estimate the C:N ratio (mol:mol, E). Particulate organic nitrogen (PON_oxi_, fmol N cell^−1^, B) and particulate organic phosphorus (POP, fmol P cell, D) were used to estimate the N:P ratio (F). PON_c_ and PON_oxi_ refer to the complete combustion method and the wet oxidation method, respectively (see methods). Each content was normalized by the cell abundance estimated at the same time point. Time on the X axis is expressed in days, starting from the beginning of the high-frequency monitoring phase, five days after the transfer to a P_i_-depleted medium. The dotted lines represent sampling in the first culture triplicate and continuous lines represent samples taken in the second triplicate (see methods). The red arrow indicates the time of DOP addition. The dashed horizontal line represents the Redfield ratio. White and grey shades represent light and dark periods, respectively.

During the recovery phase, the cellular carbon content fluctuated within a similar range as during the P_i_-depleted phase (**Figure 7A**). In contrast, the cellular N content showed much wider and more regular oscillations, between a minimum level equivalent to that of the P_i_-depleted phase, and a 50% higher maximum (29 to 63 fmol N cell^−1^; **Figure 7D**). As a result, the C:N ratio in cells followed marked, diel oscillations around an average value of 8.8 (**Figure 7E**), which is lower than the average observed at the end of the P_i_-depleted phase but still above the Redfield value of 6.6. These fluctuations point to an increase in the carbon and nitrogen fixation efficiencies compared to the P_i_-depleted phase. Likewise, the particulate P content shows notably higher levels in the recovery phase, with a 2.4-fold increase (**Figure 7D**). Considering the first point of replicate A as an outlier, the average cellular P content was 2.6 ± 0.4 fmol P cell^−1^ in the recovery phase, against 1.1 ± 0.2 in the P_i_-depleted phase. These results corroborate the strong decrease in DOP concentrations and point to an efficient DOP hydrolysis and incorporation of the released P_i_ in cells. The corresponding N:P ratio thus dropped to values in the range of 10.3 to 15.5, below the Redfield ratio, with an average of 13.0 ± 1.6 (n = 24) (**Figure 7F**). Available records of the N:P composition in *C. watsonii* under nutrient-replete conditions indicate a possible wide fluctuation of the cellular P content, with reports of N:P ratios (mol:mol) from less than 5 (51) to over 30 (52). In a previous study conducted in P replete conditions with P provided either as P_i_ or DOP, N:P ratios in the same *C. watsonii* strain ranged between 7.9 and 9.8 (14). With over twofold higher N:P ratios observed in P_i_-depleted cells in the present study, the cell stoichiometry was imbalanced and shifted. The prolonged P_i_ deficiency resulted in a far lower relative P content than P-replete conditions and we suspect cells had reached their minimum P quota.

## Discussion

The impact of nutrient limitation on nitrogen fixation in the open ocean is a widely studied topic, given the significance of diazotrophs in the input of new nitrogen in the system. The current literature provides several studies highlighting the role of P_i_ in the growth and nitrogen fixation rates of filamentous cyanobacteria, such as *Trichodesmium* (53) (54) (55) (56) (57). However, our understanding of how P_i_ influences the physiology of unicellular photosynthetic diazotrophs, especially those in the *Crocosphaera* genus, remains more limited, as does our knowledge of how P_i_ availability might affect their growth. To address these questions, we conducted a multidisciplinary study that included the functional analysis of six *Crocosphaera* genomes from three different species (*watsonii*, *chwakensis*, and *subtropica*). Additionally, we analyzed the transcription of a wide range of genes involved in the response to P_i_ stress in *C. watsonii*. Last, we monitored the growth of this cyanobacterium under P_i_-depletion, tracked the changes in its C, N and P composition, and assessed its ability to use alternative sources of P.

The estimated bacterial contamination of 3 to 7 % of the total biomass is rather low in such long-term cultures; we can reasonably consider that the changes in the C:N:P composition of the total biomass reflects that of *Crocosphaera*. Besides, from a biogeochemical point of view, all mass fluxes eventually derive from *Crocosphaera*; the medium did not contain any source of N (apart from the vitamins) so the contaminating bacteria can only feed on *Crocosphaera* C and N exudates. Also, due to the specificity of the genetic analysis, all gene expressions reported here exclusively relate to *Crocosphaera*.

We identified genes involved in P_i_ import, DOP utilization, perception of P_i_ limitation, and regulation of P_i_ stimulus expression in the six analyzed genomes, noting some variations in the composition of these genes (**Figure 1, Supporting information Table S2**). Regarding import, all six strains contained a complete high-affinity P_i_ uptake system. Our analysis refined the identification of these genes, correcting ambiguities caused by annotation errors. Notably, the *pstA* gene, which encodes one of the two membrane subunits of this system, is absent from the *C. watsonii* genome and is considered a pseudogene in NCBI. However, the literature previously listed two copies of the gene in this strain [18]. Through tBlastn analysis, we demonstrated that said *pstA* gene was mistakenly annotated as two separate small fragments in the NCBI database but is actually a unique gene. We conclude that the eight *Crocosphaera* strains have a similar PstS ABC transporter (**Figure 2**). The low-affinity P_i_ permease PitA, whose encoding gene was identified in *C. watsonii WH8501*, *C. watsonii* WH0002, and *C. watsonii* WH0401, was also found in the protein dataset of four other *C. watsonii* strains (*C. watsonii* WH8502, *C. watsonii* WH0402, *C. watsonii* WH0003, and *C. watsonii* WH0005), suggesting it is well conserved in the *C. watsonii* representatives of the *Crocosphaera* genus. In contrast, no *pitA* ortholog was found in the genomes of *C. subtropica* and *C. chwakensis*. Should PitA serve to facilitate P_i_ entry in *Crocosphaera*, this activity might have been acquired by, or maintained in, the *C. watsonii* strains only (**Figure 1**).

Genes encoding a complete phosphite and phosphonate transport system are absent in the genomes of *C. watsonii* strains, but are present in *C. subtropica* and *C. chwakensis*, indicating a possible adaptive speciation event. Given that phosphonates are produced, consumed and therefore found in all marine waters, including oligotrophic regions (58), we wonder what environmental conditions may have driven this divergence in their genomic content (**Figure 1**, **Figure 2, Supporting information Table S2**). This is all the more puzzling as an inverse relationship was highlighted between phosphonate catabolic genes and P_i_ availability across a range of marine basins, suggesting that phosphonates are a source of phosphorus for the bacterioplankton in oligotrophic areas (59). Future metagenomic surveys will be necessary to determine whether the genes involved in phosphonate uptake and degradation are expressed and functionally active in *C. subtropica* and *C. chwakensis*.

The analysis of the genes involved in DOP hydrolysis underscores the accuracy of domain-search approach to achieve a rigorous functional annotation. For instance, a gene annotated as *phoA* in the *C. watsonii* WH8501 genome suggested the presence of an alkaline phosphatase (15). However, we did not find a *phoA* gene in this strain genome; our results reveal instead that said gene encodes a 5’ nucleotidase. Conversely, a *phoX* gene is present and conserved in all the *Crocosphaera* genomes studied. At least one other gene encoding an alkaline phosphatase of another class (*phoA*, *phoD*, p*hoV*, *dedA*) was also found for each genome. To verify that the absence of a specific gene is not an artifact of our methodology, we additionally performed a tBlastN analysis for these missing genes. Apart from the *pstA* gene previously mentioned, our results confirmed the absence of all the genes that we did not find using the domain approach (**Figure 1).**

Our data collectively suggest that genes related to DOP hydrolysis are present in *Crocosphaera* genomes but raise the question as for which enzyme(s) are actually operating. The next challenge is to associate each gene with a specific DOP-scavenging activity. Biochemical assays using the MUF-P reagent have demonstrated DOP hydrolytic activities in cultures of *C. watsonii* grown with DOP supplementation (14) (17). Similarly, we observed comparable activities when P_i_-depleted cultures were shifted to the DOP-recovery phase (**Table 1**). Although *phoX* transcript levels were low, we cannot dismiss the possibility that the measured activity might be attributed to the PhoX enzyme. Under our experimental conditions, *phoX* expression might have reached a steady-state level following the phase of P_i_ depletion. Also, the transcription of genes encoding hydrolytic activities may be constrained in time. In their experiment, Pereira and colleagues (17) observed a notable increase in APA in cultures upon transfer to a P_i_-depleted medium, which confirms that APA responds to P_i_ stress. In contrast, APA was low in our cultures, but these measurements were taken after 5 days of P_i_ starvation. In parallel, the monitored C, N, P cellular composition points to a strongly skewed cell stoichiometry, with low N contents and very low, possibly minimal, P content (**Figure 7**). The synthesis of hydrolytic enzymes was therefore likely impaired by the lack of nutrients and cellular reserves. We suspect that the prolonged stress conditions had brought cultures in such a P-depleted state that cells could no longer devote sufficient energy (ATP) and possibly also nitrogen to sustaining an active synthesis of hydrolytic enzymes. But as we added DOP, APA was soon restored. We postulate that the APA level that we measured at the end of the P_i_-depleted phase, although much lower than that we measured three days after the addition of DOP, was not null. Some enzymes may also have remained active for a few days. This activity must have been sufficient for cells to start acquiring P again, which re-launched both the P and N metabolism, as demonstrated by the wider amplitude of the cellular N and P contents (**Figure 7**), allowing cells to devote again nutrients to the synthesis of hydrolytic enzymes.

If not due to alkaline phosphatases, the hydrolyzing activity of DOPs in *Crocosphaera* might be attributed to enzymes such as nucleotidases or glycerol phosphatases, should these be able to cleave MUF-P. The genes encoding these enzymes were transcribed in our experiments, but their sequences lack periplasmic targeting sequences (**Supporting information Table S3**). Given that the MUF-P reagent does not penetrate cells and only reacts with periplasmic or extracellular enzymes (60), we conclude that the identified nucleotidases or glycerol phosphatases are unlikely to be the source of the measured APA activity in healthy cells, whose membrane is intact. However, we cannot completely rule out the fact that, in such stressful conditions, in particular after days of P_i_ starvation, part of the population may have presented altered membranes or entered apoptosis. If the cellular content of some cells was released into the environment, then the APA assay might in part reflect the activity of nucleotidases or glycerol phosphatases.

Recently, a novel alkaline phosphatase enzyme (called Psip), which shares low sequence similarity with the PhoA, D, V, and X enzymes, was identified in some *Prochlorococcus* and *Synechococcus* strains (47). This enzyme had initially been designated Psip1 (for Phosphate-induced protein 1 (30). We discovered an ortholog of *psip1* exclusively in the genome of *C. chwakensis*. Therefore, the related enzyme cannot either be responsible for the DOP hydrolytic activity observed in our experiment on *C. watsonii* WH8501. Note that the name Psip is also used in the literature for a transcriptional regulator involved in P_i_ stress signaling in the annotation of *C. watsonii* genome (**Figure 1**). The multiple use of a same name for an enzyme and a regulator, as well as annotation errors (as pointed out above for *phoA*), are the source of biases and confusion in the current databases. This underscores the need for improved annotation of P_i_-related genes in cyanobacterial genomes.

In *C. watsonii* cultures, the transcription of genes involved in P_i_ uptake (*som, sphX, pstS, ugpC*), signaling (*ptrA*), and DOP utilization (5’nucleotidase, *ugpQ*, *phoX*) during a period of P_i_ limitation, and in response to DOP addition (**Figure 4, Supporting information Figure S1)**, demonstrates that the induction of these genes mediates the adaptive response to this stress. This response also resulted in changes in physiological parameters, such as cell size and elemental C, N, P contents (**Figures 6**, **7**). The survival of the cultures throughout the long, P_i_ starvation period as indicated by the relatively stable cell abundance, higher average C:N ratio and collapsed N:P ratio, reflects a certain metabolic resilience facing P_i_ depletion over several generation times (**Figure 6**, **Table 1**). The pseudo-cyclic increases and decreases in cell volume, C, N, P contents and skewed C:N and N:P ratios observed at the end of the P_i_-depleted phase altogether indicate that under phosphorus starvation, cells managed to maintain a basal activity but showed a relative inability to divide. If some cell division occurred, it could at most compensate for cell decay (**Figure 7**, **Table 1**). All the measured physiological parameters indicate that cells maintained their viability during the limitation period. The restored diel cyclicity and cells C, N, P contents at the end of the experiment reinforce this conclusion, demonstrating a recovery of growth supported by the DOP addition (**Figures 6**, **7**). These findings reveal the metabolic resilience of *C. watsonii* and underscore the profound impact of P_i_ stress on both energetic processes and nitrogen metabolism. Additionally, they illustrate the crucial role of P_i_ stress-response genes in this adaptive strategy.

## Contributions

Conceptualization: SR, AL. Investigation: ET, CC, SR, AL, OC, BM, EOR, YD, MPP. Formal analysis: ET, AL, SR, CC, VD, YD. Validation: AL, ET, SR, YD. Visualization: CC, ET, AL, SR. Supervision: SR, AL, EOR. Paper writing: AL, SR. Funding acquisition: SR.

## Acknowledgments

This study was supported by the LEFE-CYBER (CNRS) program. Work in AL lab was funded by Agence Nationale pour la Recherche Scientifique” (ANR-21-CE20-0025-01). Chloé Caille was funded by a doctoral fellowship from ED129 at Sorbonne University. Flow cytometry analyses were conducted at the SU/CNRS BioPIC Imaging and Cytometry platform of Banyuls Oceanologic Observatory under the technical support of EMBRC-France. We are grateful to the Bio2Mar platform (http://bio2mar.obs-banyuls.fr) for providing access to the Victor 3 plate reader. The authors thank Stéphanie Champ for assistance with the RNA extraction procedure. Part of the bioinformatics analyses were performed on the Core Cluster of the Institut Français de Bioinformatique (IFB) (ANR-11-INBS-0013).

## Appendix

### Appendix Supplementary Tables

**Appendix Table S1.**
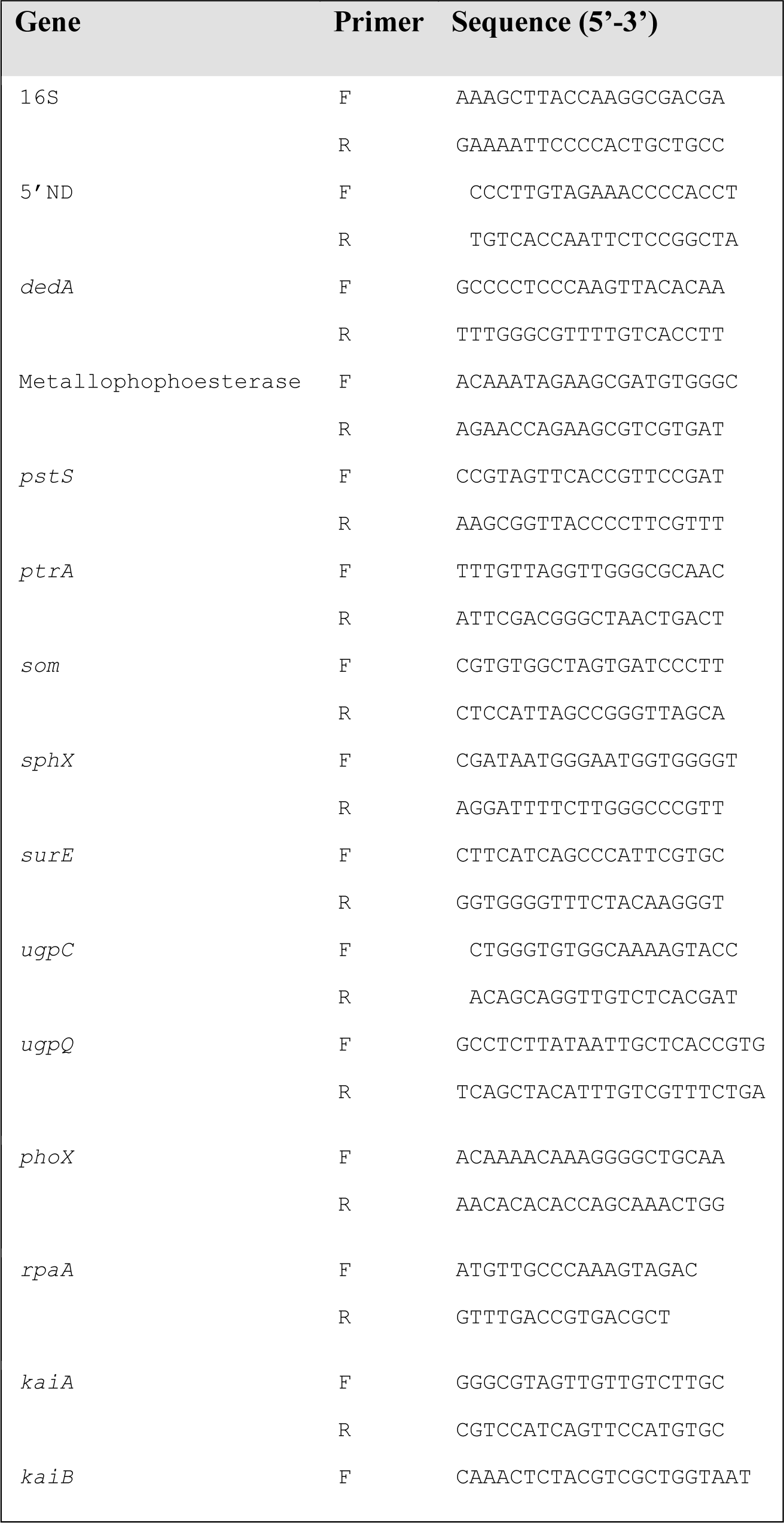
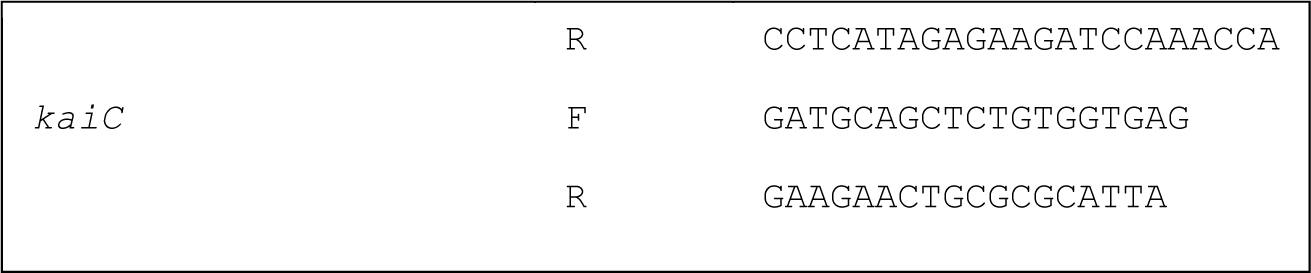
Primer sequences and amplification conditions for the genes used in RT-qPCR.

**Appendix Table S2 (Excell file):**

**Sheet 1. List of *Crocosphaera* genomes used in this study.** The cell size features were obtained from the literature.

**Sheet 2. Protein seeds (or reference proteins) used in this study.** Accession numbers (from Uniprot www.uniprot.org and Refseq www.ncbi.nlm.nih.gov databases) are indicated as well as the targeted molecules and the functional domain patterns (see **Experimental procedures**). The targeted molecule associated to each seed was obtained from the literature. Within the domain patterns, domains are shown in the order of appearance in the protein and separated with a star (*). Domain abbreviations and their functional descriptions of the domains are shown in **Sheet 3**. Other data were retrieved from the Uniprot or NCBI databases. Similar or identical seeds (from distinct organisms) are grouped into one seed (*e.g.* GppA-PpX).

**Sheet 3. List of functional seed domains Sheet 4. List of protein orthologs.**

**Sheet 5. Distribution of protein orthologs in *Crocosphaera* species**

**Sheet 6. Distribution of protein homologs in group of *Crocosphaera* species**

**Appendix Table S3:** Prediction of cellular location of DOP-hydrolyzing enzymes in *C. watsonii* WH8501. The presence of signal peptides Sec or Tat was analyzed using Signal IP-6.01 software (https://services.healthtech.dtu.dk/services/SignalP-6.0/)

| Protein | Signal peptide | Type/position (amino acids) | Predicted cellular location |
| --- | --- | --- | --- |
| Alcaline phosphatase PhoX | Yes | TAT/ 1-24 | Extra-cytosolic |
| Phosphatase like DedA | No |  | Cytosolic |
| 5'Nucleotidase SurE | No |  | Cytosolic |
| Metallophosphoesterase | No |  | Cytosolic |
| Glycerolphosphoesterase UgpQ | No |  | Cytosolic |

### Appendix Supplementary Figures

**Appendix Figure S1:**
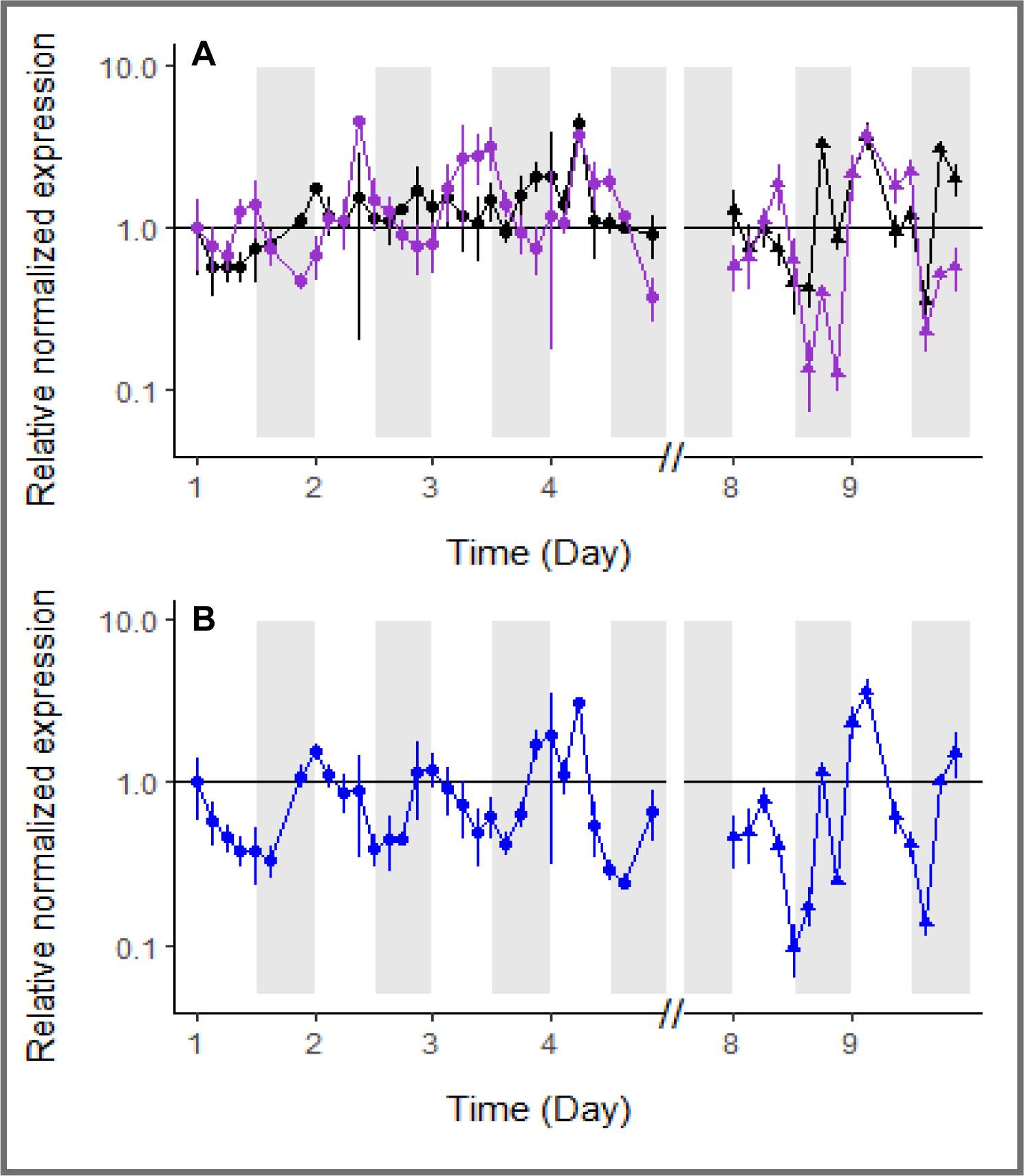
Relative normalized gene expression of *C. watsonii* during the P_i_-depleted phase (closed circles, left panels) and the DOP-recovery phase (closed triangles, right panels). RT-qPCR of the genes encoding a potential 5’-nucleotidase, *surE* (in black, **A**), a metallophosphoesterase (in purple), and the alkaline phosphatase-like, *dedA* (in blue, **B**). The Y-axis is logarithmic. All points are normalized by the 16S expression at the same point and relative to the first sampling point (Day 1, L0). Expression variability of biological duplicates and analytical duplicates are represented by error bars. White and grey shades represent the light and dark periods, respectively; red arrows indicate the day of DOP addition.

**Appendix Figure S2:**
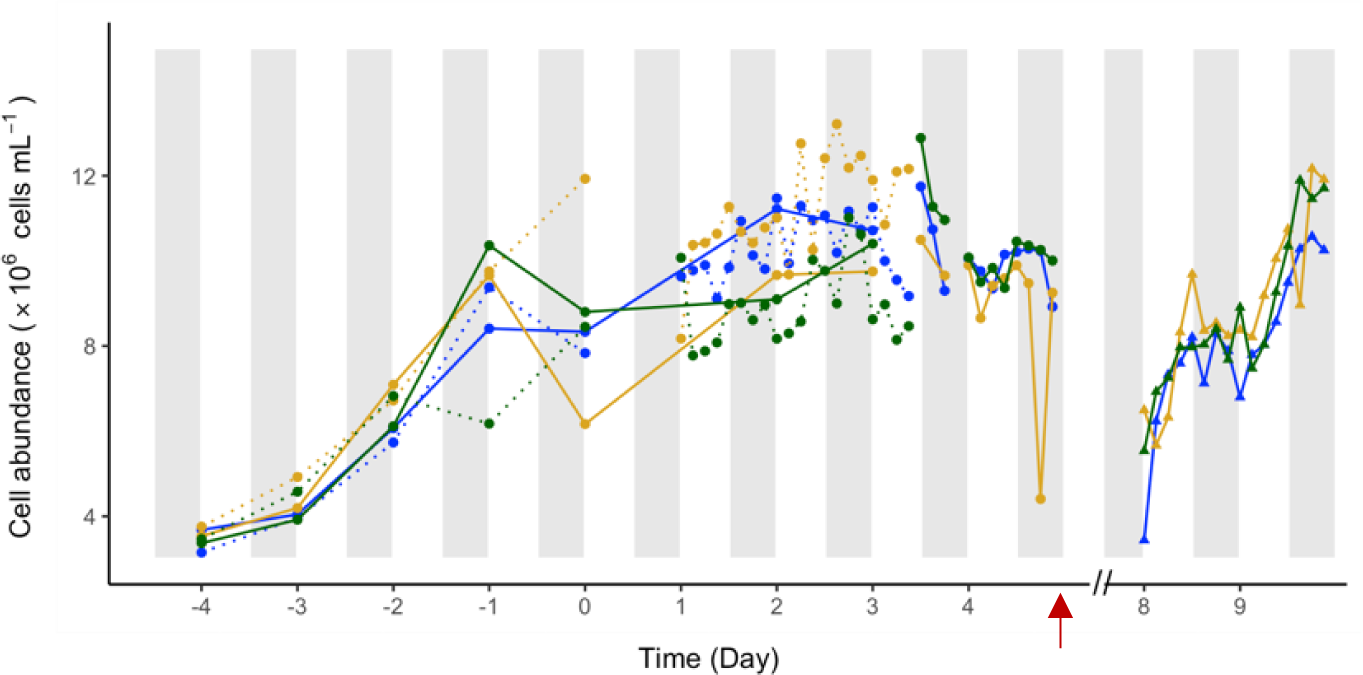
Diel fluctuations in *C. watsonii* total population cell abundance (cell.mL^−1^) measured in each triplicate (blue, yellow and green) over the entire P_i_-depleted period. Cultures were transferred to the P_i_-depleted medium on Day −4 and the high-frequency monitoring started on Day 1. The dotted lines represented sampling in the first triplicate and solid lines samples taken in the second triplicate. The red arrow marks the day of DOP addition and samples of the DOP-recovery phase were realized Days 8 and 9.

**Appendix Figure S3:**
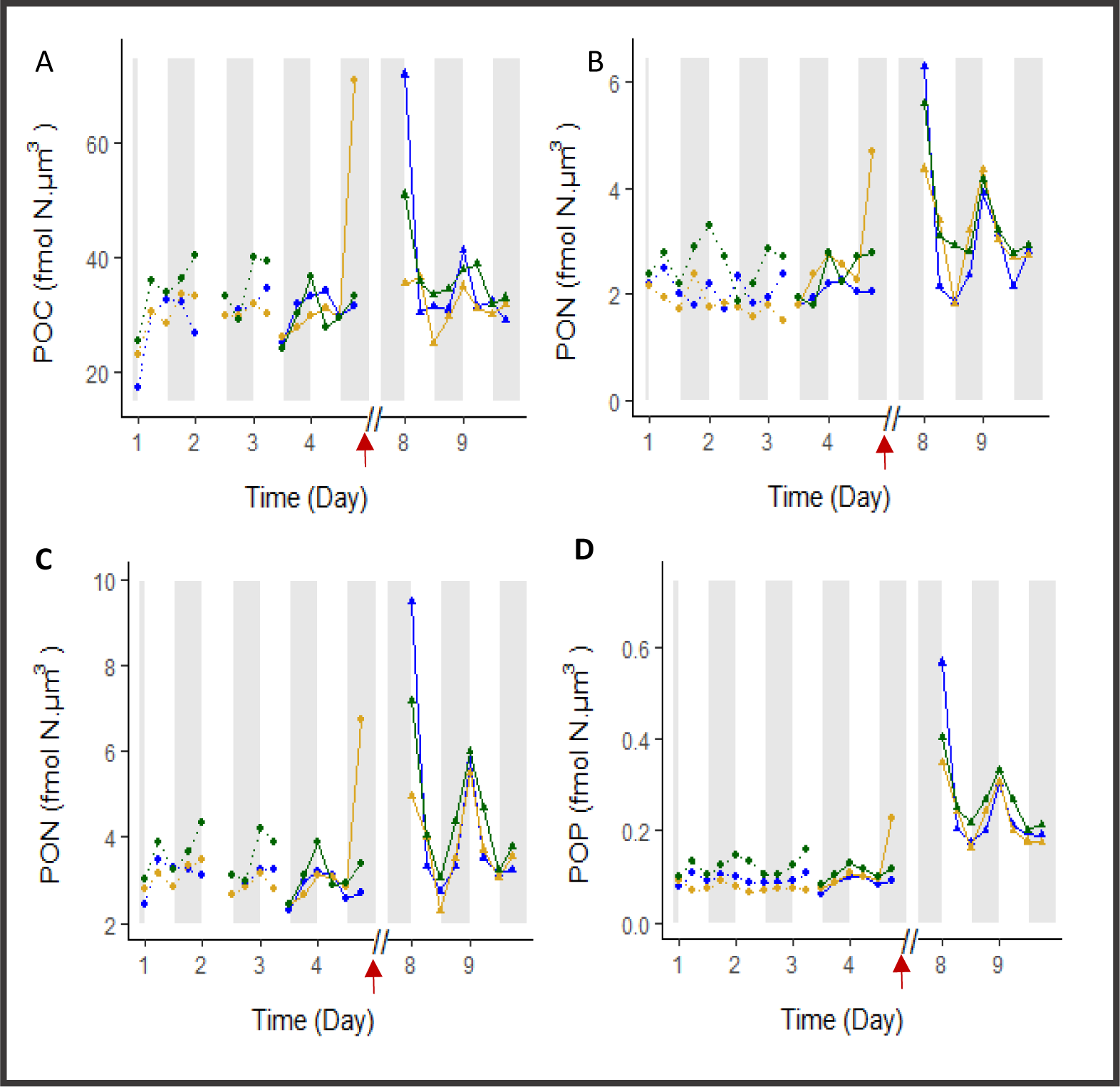
Diel fluctuations of *C. watsonii* C, N and P cell contents in each replicate (blue, yellow and green) during the P_i_-depleted (Day 1 to Day 4) and DOP-recovery (Day 8 and Day 9) phases. Particulate organic carbon (POC, fmol C µm^3^, A) and particulate organic nitrogen (PON_c_, fmol N µm^3^, C) were used to estimate the C:N ratio (mol:mol, E). Particulate organic nitrogen (PON_oxi_, fmol N µm^3^, B) and particulate organic phosphorus (POP, fmol P µm^3^, D) were used to estimate the N:P ratio (F). PON_c_ and PON_oxi_ refer to the complete combustion method and the wet oxidation method, respectively (see methods). Each content was normalized by the biovolume (µm^3^) estimated at the same time point. Time on the X axis is expressed in days, starting from the beginning of the high-frequency monitoring phase, five days after the transfer to a P_i_-depleted medium. The dotted lines represent sampling in the first culture triplicate and continuous lines represent samples taken in the second triplicate (see methods). The red arrow indicates the time of DOP addition. The dashed horizontal line represents the Redfield ratio. White and grey shades represent light and dark periods, respectively.

