## Supplemental Table 1 for "Unveiling *Crocosphaera* responses to phosphorus depletion: insights from genome analysis and functional characterization"

### Supplementary Tables

**Appendix Table S1.** Primer sequences and amplification conditions for the genes used in RT-qPCR

| Gene | Primer | Sequence (5’-3’) |
|---|---|---|
| 16S | F | AAAGCTTACCAAGGCGACGA |
| | R | GAAAATTCCCCACTGCTGCC |
| 5’ND | F | CCCTTGTAGAAACCCCACCT |
| | R | TGTCACCAATTCTCCGGCTA |
| *dedA* | F | GCCCCTCCCAAGTTACACAA |
| | R | TTTGGGCGTTTTGTCACCTT |
| Metallophophoesterase | F | ACAAATAGAAGCGATGTGGGC |
| | R | AGAACCAGAAGCGTCGTGAT |
| *pstS* | F | CCGTAGTTCACCGTTCCGAT |
| | R | AAGCGGTTACCCCTTCGTTT |
| *ptrA* | F | TTTGTTAGGTTGGGCGCAAC |
| | R | ATTCGACGGGCTAACTGACT |
| *som* | F | CGTGTGGCTAGTGATCCCTT |
| | R | CTCCATTAGCCGGGTTAGCA |
| *sphX* | F | CGATAATGGGAATGGTGGGGT |
| | R | AGGATTTTCTTGGGCCCGTT |
| *surE* | F | CTTCATCAGCCCATTCGTGC |
| | R | GGTGGGGTTTCTACAAGGGT |
| *ugpC* | F | CTGGGTGTGGCAAAAGTACC |
| | R | ACAGCAGGTTGTCTCACGAT |
| *ugpQ* | F | GCCTCTTATAATTGCTCACCGTG |
| | R | TCAGCTACATTTGTCGTTTCTGA |
| *phoX* | F | ACAAAACAAAGGGGCTGCAA |
| | R | AACACACACCAGCAAACTGG |
| *rpaA* | F | ATGTTGCCCAAAGTAGAC |
| | R | GTTTGACCGTGACGCT |
| *kaiA* | F | GGGCGTAGTTGTTGTCTTGC |
| | R | CGTCCATCAGTTCCATGTGC |
| *kaiB* | F | CAAACTCTACGTCGCTGGTAAT |
| | R | CCTCATAGAGAAGATCCAAACCA |
| *kaiC* | F | GATGCAGCTCTGTGGTGAG |

| | | |
|---|---|---|
| | R | GAAGAACTGCGCGCATTA |
